# Importin α4 deficiency induces psychiatric disorder-related behavioral deficits and neuroinflammation in mice

**DOI:** 10.1101/2024.04.18.590002

**Authors:** Koki Sakurai, Makiko Morita, Yoshiatsu Aomine, Mitsunobu Matsumoto, Tetsuji Moriyama, Emiko Kasahara, Atsuo Sekiyama, Mayumi Otani, Rieko Oshima, Kate L. Loveland, Masami Yamada, Yoshihiro Yoneda, Masahiro Oka, Takatoshi Hikida, Yoichi Miyamoto

**Author notes:** Corresponding Authors: Masahiro Oka, Department of Regulation of Infectious Cancer, Research Institute of Microbial Diseases (RIMD), Osaka University, 3-1 Yamadaoka, Suita, Osaka, 565-0871, Japan.; Takatoshi Hikida, Laboratory for Advanced Brain Functions, Institute for Protein Research, Osaka University, 3-2 Yamadaoka, Suita, Osaka, Japan; Yoichi Miyamoto, Laboratory of Biofunctional Molecular Medicine, National Institutes of Biomedical Innovation, Health and Nutrition, Saito-asagi 7-6-8, Ibaraki, 567-0085 Osaka, Japan.

## Abstract

Importin α4, which is encoded by the *Kpna4* gene, is a well characterized nuclear-cytoplasmic transport factor known to mediate transport of transcription factors including NF-κB. Here, we report that *Kpna4* knock-out (KO) mice exhibit psychiatric disorder-related behavioral abnormalities such as anxiety-related behaviors, deceased social interaction and sensorimotor gating deficits. Contrary to a previous study predicting attenuated NF-κB activity as a result of *Kpna4* deficiency, we observed a significant increase in expression levels of NF-κB genes and pro-inflammatory cytokines such as *TNFα*, *Il1β* or *Il-6* in the Prefrontal Cortex or Basolateral Amygdala of the KO mice. Moreover, examination of inflammatory responses in primary cells revealed that *Kpna4* deficient cells have an increased inflammatory response, which was rescued by addition of not only full-length, but also a nuclear transport deficient truncation mutant of importin α4, suggesting contribution of its non-transport functions. Furthermore, RNAseq of sorted adult Microglia and Astrocytes and subsequent transcription factor analysis suggested increases in Polycomb repressor complex 2 (PRC2) activity in *Kpna4* KO cells. Taken together, importin α4 deficiency induces psychiatric disorder-related behavioral deficits in mice, along with an increased inflammatory response and possible alteration of PRC2 activity in glial cells.

## INTRODUCTION

Accumulating epidemiological evidence have identified a plethora of genetic and environmental risk factors that contribute to the pathogenesis of psychiatric disorders. Recently, members of the importin α (karyopherin α: KPNA) family such as KPNA1 (human importin α5), KPNA3 (human importin α4), and KPNA4 (human importin α3) have been identified as possible genetic risk factors to several different psychiatric disorders including schizophrenia, depression, and substance use disorders [1–6]. These 3 importin α subtypes are expressed in the central nervous system (CNS) of humans as well as mice[7], and constitutive depletion in mice has been associated to disorder-related behaviors: *Kpna1* deficiency causing reduced anxiety and other psychiatric disorder-related behaviors[8–10], and *Kpna3* deficiency causing deficits in reward-seeking behavior[11]. In particular, a postmortem study has implicated human importin α3 (KPNA4)[12] in the pathology of schizophrenia [6], where significantly decreased nuclear factor-kappa B (NF-κB) pathway signaling, decreased p65 protein levels and nuclear activation, and *KPNA4* downregulation was found in schizophrenia brains, suggesting that decreased KPNA4 leads to deficient nuclear transport of p65 in schizophrenia patients. Moreover, in the same study, an allele in a *KPNA4* expression quantitative trait locus (eQTL) was associated to increased risk for schizophrenia, decreased *KPNA4* expression, and decreased prepulse inhibition (PPI), suggesting that *KPNA4* depletion could have roles in the pathogenesis of schizophrenia. Despite such evidence, there has been little insight on the causal relationship between *Kpna4* deficiency in relation to psychiatric disorder-related behavior.

Importin αs are a structural and functional subcategory of the importin (karyopherin) superfamily which mediate signal-dependent protein transport across the nuclear envelope[13, 14]. Importin αs participate in nucleocytoplasmic transport by forming a trimeric complex together with classical nuclear localization signal (cNLS) containing cargo proteins, as well as importin β1; another member of the importin superfamily which facilitates passage through the nuclear pore complex (NPC)[15]. Importin α subtypes show differential expression patterns in various tissues, as well as having distinct, yet “partially redundant” binding specificities[16], implying that differential importin α expression can regulate the accessibility of nuclear proteins to the nucleus[16–18]. Additionally, recent accumulating evidence suggests that importin αs are involved in non-transport functions such as spindle assembly, nuclear envelope assembly, Lamin formation, protein degradation, and chromatin alteration[15, 19–22], as well as neuron-specific functions such as axonal transport[23]; however, the physiological implications of such widespread functions are still under extensive examination.

Mouse importin α4 categorizes in the same α3 subfamily with the closely related subtype importin α3, which is encoded by the *Kpna3* gene and shares common characteristics such as cargo specificity for proteins such as Regulator of chromosome condensation 1 (RCC1), tumor suppressor p53, and Methyl-CpG binding protein 2 (MeCP2)[16, 24]. In particular, the α3 subfamily has been well characterized in the tumor necrosis factor alpha (TNF-α) induced nuclear translocation of NF-κB subunits p65 (RelA) and p50[25–28], the pathway which the previous study has suggested to be downregulated in schizophrenia patients[6]. In relation, a recent study has reported that *Kpna4* deficiency hinders NF-κB nuclear translocation in lung cells, disrupting antiviral responses against influenza resulting in higher lethality in mice[29]. Although a constitutively *Kpna4* deficient mice line has been reported to show decreased pain responsiveness and impairment of c-fos nuclear import in sensory neurons[30], there has been little investigation into the effects of *Kpna4* deficiency in psychiatric disorder-related behaviors and regulation of neuroinflammation. Further examination of such psychiatric disorder-related behaviors in *Kpna4* deficient animals is necessary to elucidate the roles of the importin α4 in regulation of brain function and behavior.

In this study, we used a recently developed importin α4 (*Kpna4*) knockout (KO) mouse line which show no apparent deficits in gross morphology, but exhibit male subfertility and deficiencies in sperm morphology, motility, and acrosome reaction capacity[21]. Using this knockout line, we found that *Kpna4* deficiency in mice induces psychiatric disorder-related behaviors, increased neuroinflammation, enhanced inflammatory responses in primary cultured astrocytes, as well gene expression patterns suggestive of enhanced inflammatory responses and altered Polycomb repressor complex 2 (PRC2) activity in sorted adult glial cells.

## RESULTS

### KO mice exhibit psychiatric disorder-related behaviors

To investigate the effects of constitutive *Kpna4* deficiency results on behavior, we conducted a behavioral test battery consisting of an open field test (OFT), elevated plus maze (EPM), Y-Maze, social interaction test, inhibitory avoidance (IA), and prepulse inhibition (PPI) tests to assess psychiatric disorder-associated behavior.

Locomotor activity and anxiety-like behavior were assessed in an open field test (OFT), where no significant differences were observed in novelty-induced locomotion (first 5 min) between all genotypes (Fig. 1A); however, KO mice spent significantly shorter durations of time in the center of the open field (Fig. 1B), suggesting higher levels of anxiety-like behavior. There was no significant difference between genotype in general levels of locomotion over the entire 60 min trial (Fig. 1C, Fig. S1A). Furthermore, similar to the results in the OFT, in the elevated plus maze (EPM) test KO mice showed significantly shorter durations of time in the open arms (Fig. 1D), suggesting higher levels of anxiety-like behavior in the KO mice.

**Figure 1.**
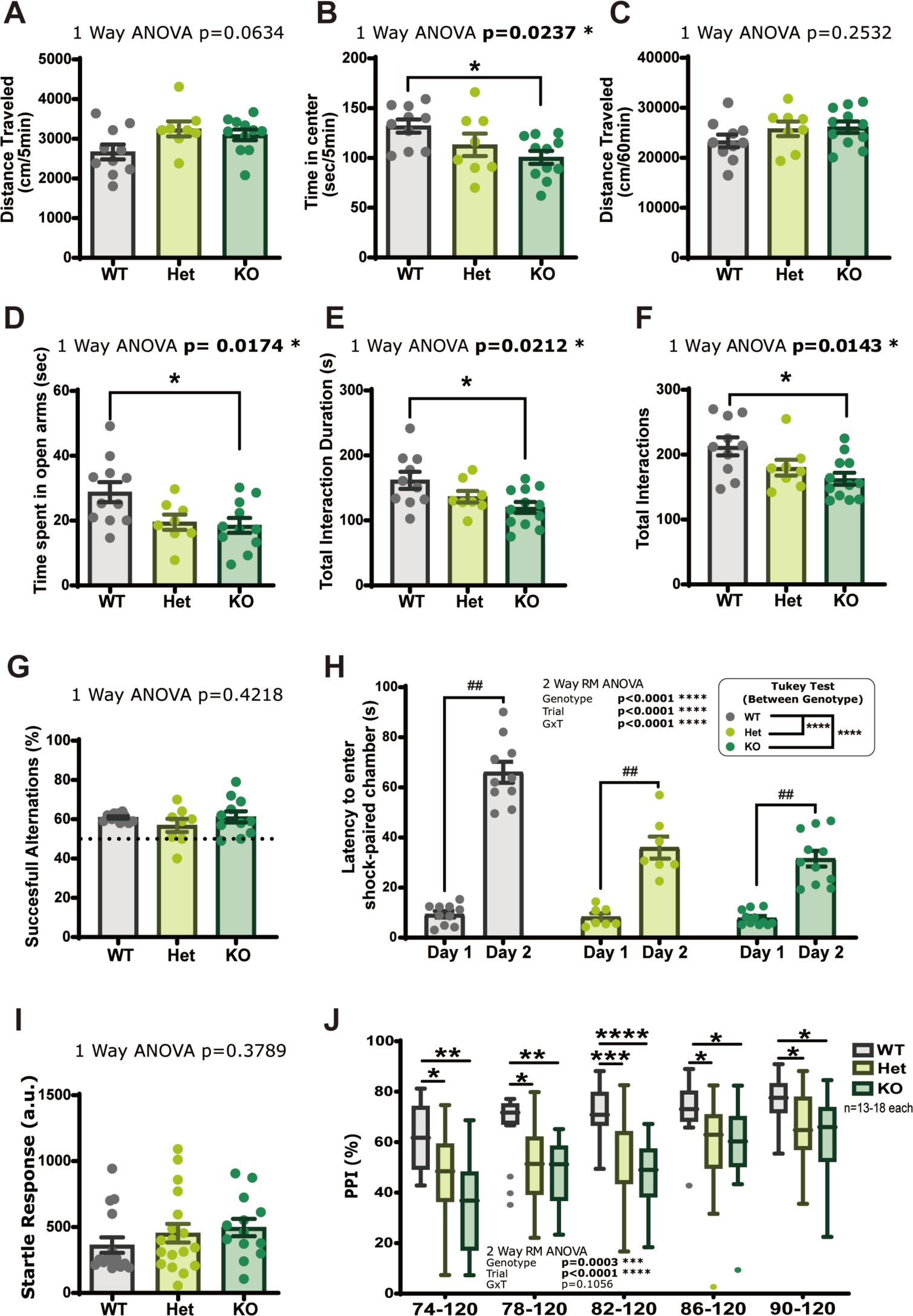
Behavioral analysis reveals psychiatric disorder-related behavioral deficits in *Kpna4* deficient mice. (A-C) OFT. The total distance (A) and percentage of the time spent in the center (B) in the first 5 min of the OFT. Total distance (C) in the entire 60 min duration of the OFT. (D) EPM. The duration of the time spent in the open arms in the EPM Test. (E-F) Social Interaction Test. The duration (E) and number (F) of total interaction behaviors in the social interaction test. (G) Percentage of successful alternations in the Y-maze test. (H) IA. Latency to step through to the dark chamber in the IA test. (I-J) PPI. (I) Startle response when presented the only startle stimulus (120 dB). (J) Level of prepulse inhibition seen in the PPI test. Bar graphs represent Mean ± SEM. Box and whisker plots represent Median (center line), First and Third Quartiles (box), ±1.5 Interquartile range (IQR) (whiskers), Data points outside ±1.5 IQR are visualized as dots. Post-hoc Tukey’s Test: ****p<0.0001, ***p<0.001, **p<0.01, *p<0.05. (A-J) WT 10-11, Het 7-8, and KO 11-12 (K-L) WT 16, Het 18, and KO 13.

We examined the social behaviors of KO mice in a reciprocal social interaction test[31], where the number and duration of contacts between the nose point of one mouse, with the nose point, body center, or tail base of the other mouse was quantified as a measure of social interaction (Total Interactions; sum of all 3 measures for each mouse). Significant decreases in KO mice were observed in both duration (Fig. 1E) and counts of social interactions (Fig. 1F), along with several individual measures (Fig. S1 B-G) in the social interaction test.

To assess if *Kpna4* depletion results in memory deficits, we assessed short-term spatial memory and avoidance learning in KO mice. In the Y-maze test, we did not observe the any significant alterations in spontaneous alternation between all genotypes (Fig. 1G). In the inhibitory avoidance task[32], 2W-ANOVA analysis revealed significant effects of both *Kpna4* deficiency (Genotype) and Test Day (Trial), as well as a significant interaction between them (Fig. 1H; 2W-RM-ANOVA; Main effect of genotype: F (2, 25) = 25.41, p<0.0001; Main effect of Test Day (Trial): F (1, 25) = 229.1, p<0.0001; Interaction: F (2, 25) = 20.96, p<0.0001). Inter-day comparison of latencies showed that WT mice showed a significantly longer latency to enter the dark chamber on day 2. In contrast, heterozygote (Het, *Kpna4^+/−^*) and KO mice showed significantly reduced latencies to enter the dark chamber compared with their WT littermates.

To test whether *Kpna4* deficiency results in sensorimotor gating deficits, we administered a PPI test against an acoustic startle stimulus. There was no significant difference in startle response to the 120dB pulse alone (Fig. 1I). In contrast, levels of PPI were significantly altered as a result of genotype (Fig. 1J; 2W-RM-ANOVA; Main effect of genotype: F (2, 44) = 9.930, p=0.0003; Main effect of Prepulse Strength (Trial): F (3.015, 132.6) = 32.05, p<0.0001; Interaction: F (8, 176) = 1.682, p=0.1056), with KO mice exhibiting significantly lower levels of PPI compared to WT mice in all types of trials (Prepluse Strength 74, 78, 82, 86, 90 dB).

### Examination of morphology and expression of Importin α subtypes in the KO brain

Examination of gross morphology in the KO brain sections revealed no apparent defects (Fig. S2A), and past studies[7] as well as examination of single cell RNAseq databases[33, 34] show that *Kpna4* expression is not region or cell type specific. Moreover, we examined if *Kpna4* deficiency results in complimentary upregulation of other importin α subtypes (*Kpna1*, *Kpna2*, *Kpna3*, *Kpna6*) in brain tissue. In the PFC, although qRT-PCR analysis showed slight but significant increase in mRNA levels of closed-related importin α3 (Fig. S2B), the importin α3 protein levels were not significantly altered (Fig. S2C, D). In addition, there was no difference in expression of other subtypes between WT and KO mice in the hippocampus (Fig. S2E). Furthermore, immunohistochemical staining showed nuclear localization of Importin α4 across several different areas in the brain (Fig. S3)

### Increased proinflammatory reactions in brains of KO mice

As Importin α4 (KPNA4) has been well characterized in the nuclear transport for NF-κB [25, 29], and its downregulation has been predicted to perturb NF-κB signaling in postmortem studies[6], we sought to examine if such perturbations occur in the brains of KO mice. Regional expression levels of NF-κB genes and proinflammatory cytokines were assessed, examining 5 regions (Prefrontal Cortex: PFC, Nucleus Accumbens: NAc, Dorsal Hippocampus: Hipp, Basolateral Amygdala: BLA, Cerebellum: Cere) sampled from mice used in behavioral testing (Fig. 2A, S4). We first quantified regional *Kpna4* mRNA levels in various regions, where expression was detected in all 5 regions with modest variation (Fig. 2B). We next quantified mRNA expression of *Rela* and *Nfkb1* in the KO brain where, unexpectedly, we observed significant upregulation of *Rela* in the PFC, as well as significant upregulation of *Nfkb1* and increasing trends of *Rela* in the in the BLA (Fig. 2C-D).

**Figure 2.**
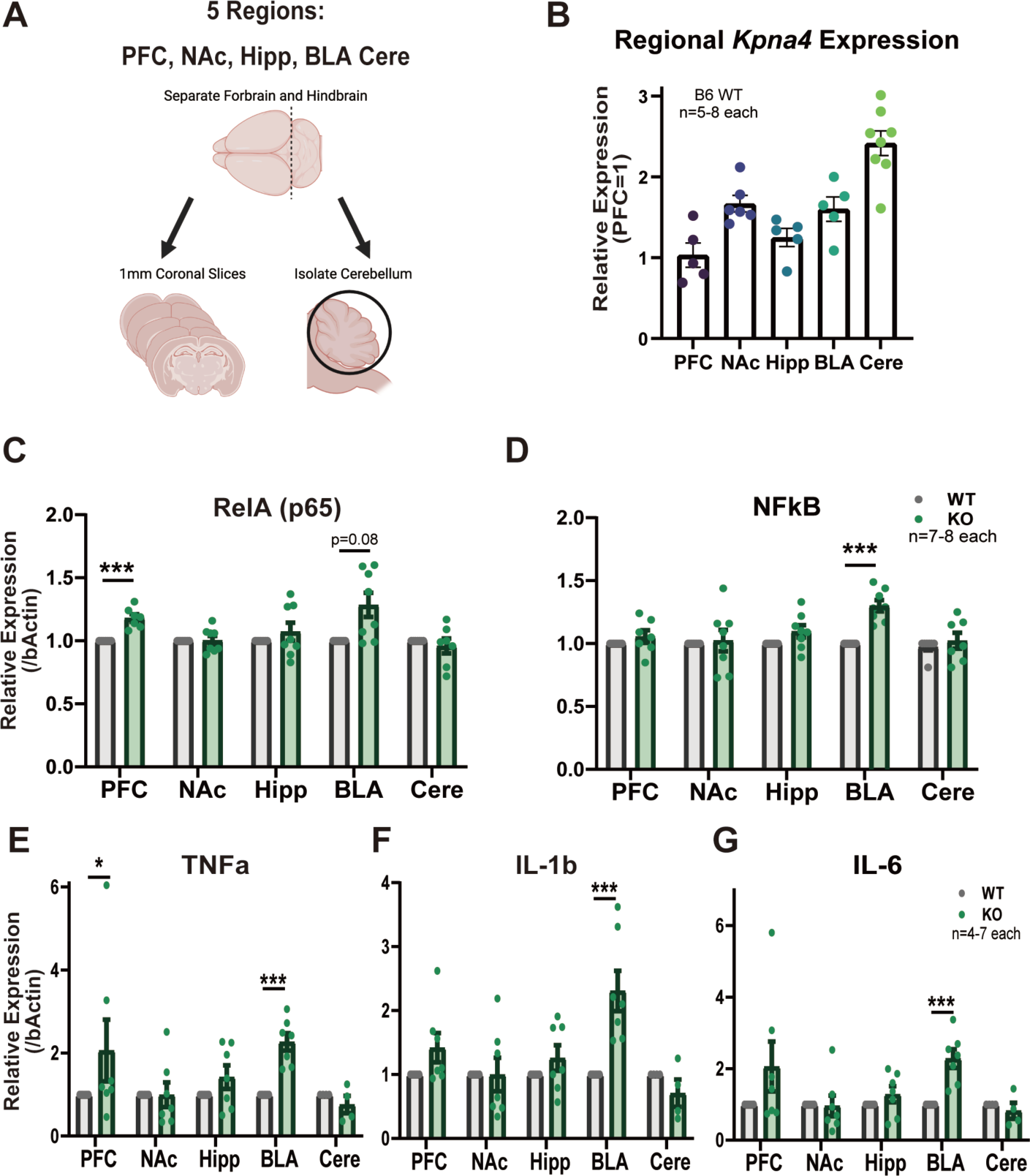
Upregulation of NF-κB genes and proinflammatory cytokines in the KO mouse brain. (A) Scheme of regional sampling (Created with BioRender.com.), coordinates shown in Fig. S3. (B) Regional expression levels of *Kpna4* (prefrontal cortex: PFC, nucleus accumbens: NAc, hippocampus: Hippo, basolateral amygdala: BLA, and cerebellum: Cere; n=6 each) (C-D) Regional mRNA expression levels (/β-Actin) of (C) *Rela* and (D) *Nfkb1* in KO and WT mice (PFC, Hippo, BLA n=5. NAc n=6. Cere n=4 each). (E-G) Regional mRNA expression levels of (E) *Tnf-α*, (I) *Il1-β*, and (G) *Il-6* in KO and WT mouse brains (PFC, NAc, Hippo, BLA n=6. Cere n=4 each). Bar graphs represent Mean ± SEM. (C-D) Mann-Whitney Test: *** p<0.001, ** p<0.01, and *p<0.05.

We further examined the regional expression of 3 typical proinflammatory cytokines: *Tnf-a*, *Il-1b*, and *Il-6* expected to be downregulated as a result of decreased NF-κB nuclear retention. Similar to *Rela* and *Nfkb1*, significant upregulation of all 3 genes was observed in the BLA of KO mice, along with significantly upregulated *Tnf-a* in the PFC, and trend towards increase of all three in the PFC and Hipp (Fig. 2E-G). Examination of systemic changes in immune-related signal proteins in the same mice did not show any significant alterations (Fig. S5). Taken together, these data suggest that KO mice have increased proinflammatory responses associated with enhanced NF-κB signaling specific to the brain.

### Increased proinflammatory activation in *Kpna4* deficient cells

As we observed such unexpected increases in NF-κB genes and proinflammatory cytokines from the KO brain, we further investigated the effects of *Kpna4* deficiency on cellular inflammatory responses using primary cultured neural cells, focusing on glial cells. We first assessed TNF-α-induced nuclear translocation of NF-κB subunit p65 in primary astrocytes (AST). Immunofluorescence analysis revealed clear nuclear translocation of endogenous p65 in response to TNF-α treatment was visible in both WT and KO cells, revealing that *Kpna4* depletion does not alter their localization to the nucleus (Fig. 3A). Notably, nuclear localization ratio of p65 in the KO cells showed significantly higher than WT (Fig. 3B). Moreover, examination the expression levels of proinflammatory cytokines revealed that *Il-1b* and *Il-6* were upregulated in KO primary AST after TNF-α stimulation (Fig. 3C-D), Taken together with the results from KO brains, our observations suggest that *Kpna4* depletion results in an increase in inflammatory activation by increasing the concentration of p65 in the nucleus, rather than inhibiting nuclear transport of p65.

**Figure 3.**
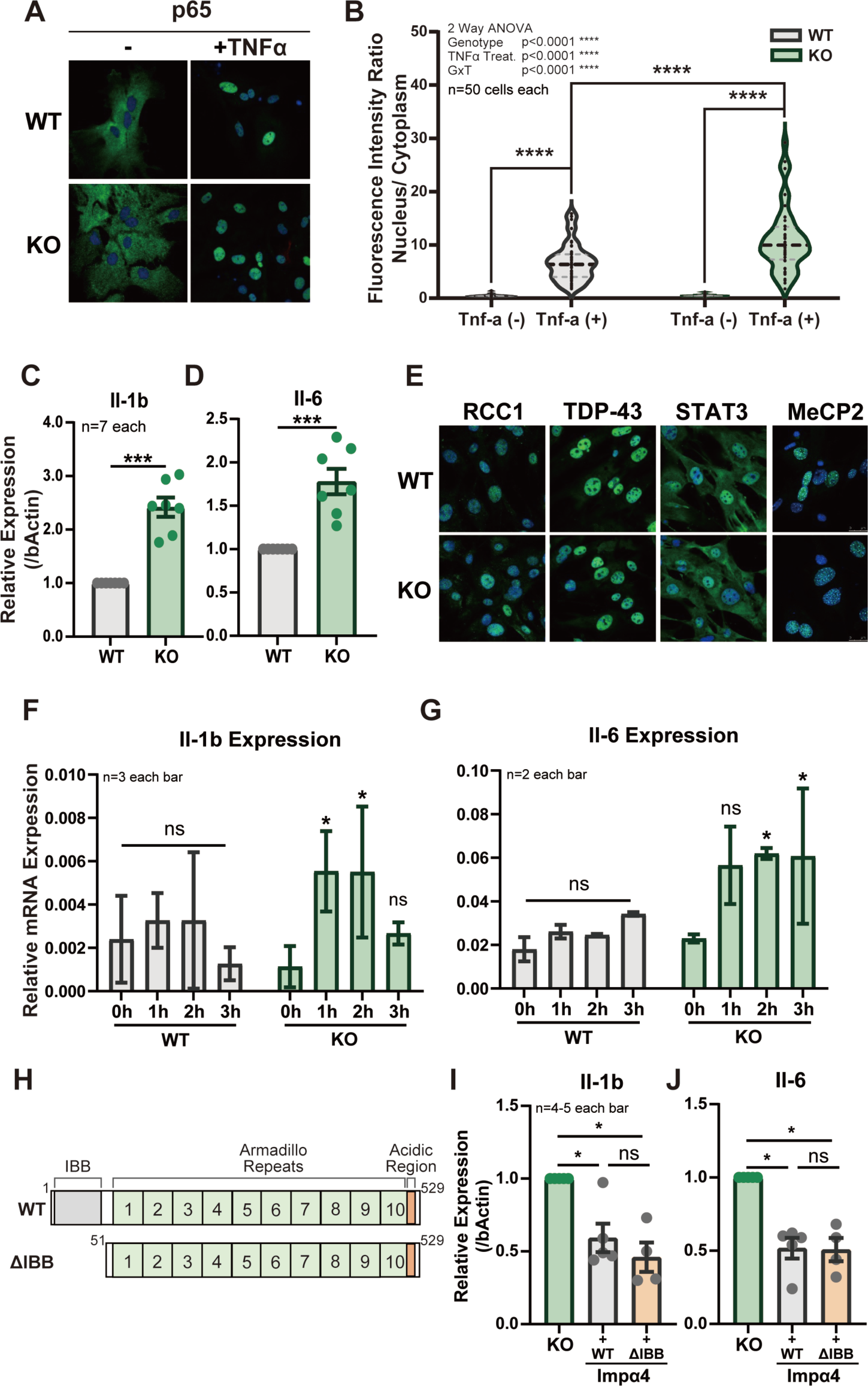
Importin α4 depletion increases proinflammatory responses in cell culture and is suppressed by reintroduction of WT or transport-deficient (ΔIBB) KPNA4. (A) Immunofluorescence analysis of p65 (RELA) in *Impα4^−/−^* Astrocytes. Nuclear translocation of p65 in response to 30 min TNF-α stimulation was observed in the primary astrocytes established from KO and WT mice. (B) Fluorescence intensity ratio of nuclear vs cytoplasmic p65 in (A) (n=50 cells each). (C-D) mRNA expression levels of (C) *Il1-β* and (D) *Il-6* (/β-Actin) in KO and WT primary astrocytes after 3 h stimulation with TNF-α (n=7 each). (E) Immunofluorescence analysis of RCC1, TDP-43, STAT3, MeCP2 in (unstimulated) KO and WT MEFs. (F-G) Time-course changes of (F) *Il1-β* and (G) *Il-6* mRNA expression in KO and WT astrocytes (n=2-3 each bar). (H) Schematic representation of importin α4 (KPNA4)-WT and ΔIBB mutant transfected to KO cells. (I-J) Expression levels of (I) *Il1-β* and (J) *Il-6* (/β-Actin) in KO MEFs transfected with EGFP-importin α4 (Imp-α4)-WT, EGFP-Imp-α4-ΔIBB, or EGFP control after 1h stimulation with TNF-α (n=4-5 each). Bar graphs represent Mean ± SEM. Violin plots represent Median (solid line), First and Third Quartiles (dotted line). (B) Post-hoc Sidak’s Test: ****P<0.0001. (C-D) Mann-Whitney Test: ***P<0.001. (F-G) Dunnett’s test (vs 0h) *P<0.05. (I-J) Dunn’s test *P<0.05.

Furthermore, in our examination MEF cells established from WT and KO mice, nuclear translocation of the other typical cargos specific for the importin α3 family such as RCC1[35, 36], TDP-43[37], STAT3[38], and MeCP2[24] were maintained in KO MEFs, indicating that *Kpna4* deficiency does not disturb their nuclear transport (Fig 3E). Similar to our results in primary AST cells, time-course monitoring of *Il-1b* and *Il-6* expression after TNF-α stimulation showed that KO MEFs show increased inflammatory activation (Fig. 3F-G).

In a previous study, we have reported that epigenetic alteration in the testis of KO mouse leads to altered gene expression, resulting in abnormal sperm formation and infertility[21]. To address whether such epigenetic functions of importin α4 are involved in the aberrant upregulation of cytokine expression, we transfected KO MEFs with importin α4 ΔIBB (IBB domain truncated, transport deficient) mutant as well as importin α4 WT to rescue the increases in cytokine expression (Fig 3H). Both EGFP-importin α4 WT and EGFP-importin α4 ΔIBB transfected cells showed significantly decreased *Il1b* and *Il-6* expression after TNF-α stimulation compared to EGFP transfected controls (Fig 3I-J). This result suggests that the effects of importin α4 in suppressing aberrant proinflammatory activation is dependent on non-transport functions such as chromatin alteration and epigenetic regulation, rather than its well characterized transport functions.

### Increased proinflammatory signaling and altered Polycomb Repressive Complex activity in KO glial cells

To investigate the molecular alterations behind the unexpected increase in proinflammatory activation induced by *Kpna4* deficiency, we examined individual glial cell types to understand gene expression profiles in response to a proinflammatory stimulus. We applied a tissue disassociation-cell sorting strategy utilizing magnetic cell sorting (MACS) or florescence activated cell sorting (FACS), to isolate both microglial (MG) and AST populations from mice administered LPS to stimulate inflammation (Fig 4A; S6). In line with our tissue and cellular experiments, we saw significant upregulation of *Il-6* expression in MG and AST isolated from adult KO mice using MACS, along with an increasing trend in *Tnf-a* and *Il-1b* (Fig S7). Thus, we proceeded to isolate these populations with FACS to examine their gene expression profiles using RNAseq. In contrast with MG and AST, similar assessment of *Tnf-a*, *Il-1b*, and *Il-6* in non-neural immune cells (peritoneal macrophages) collected from the same mice showed no significant difference in expression of typical proinflammatory cytokines (Fig S8).

**Figure 4.**
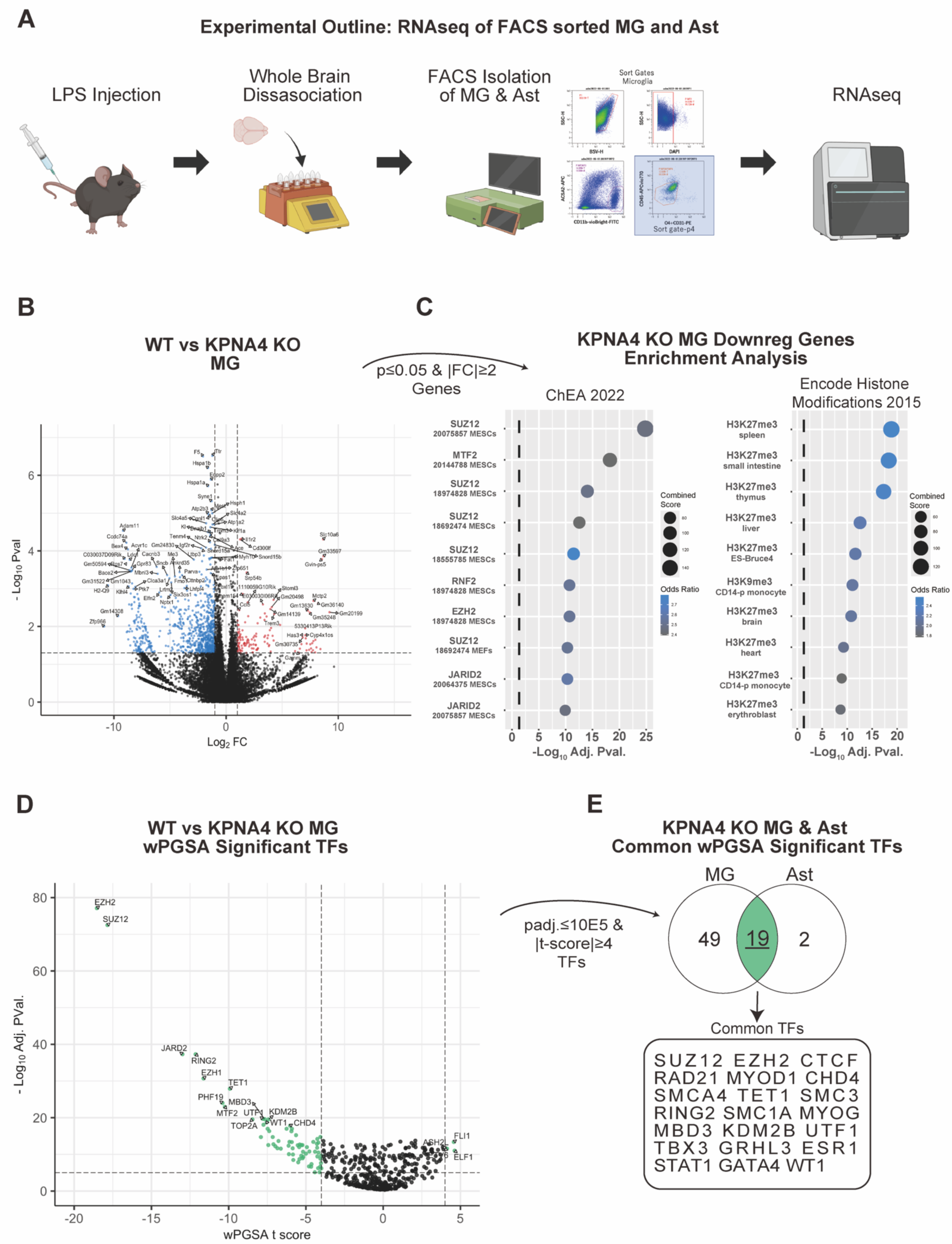
RNAseq analysis of KO microglia reveal alterations genes regulated by Polycomb Repressor Complex. (A) Outline of cell sorting experiments from KO and WT mice (Created with BioRender.com.). (B) Volcano plot of genes with altered expression in KO MGs in comparison to WT mice. Horizontal axis indicates log_2_ Fold Change, and vertical axis indicates -log_10_ p value. Colored points indicate genes with p ≤ 0.05 and |Fold Change| ≥2. (C) Results of enrichment analysis by Enrichr on MG downregulated gene list. Dotted line indicates adjusted P=0.05. (D) Volcano plot of wPGSA analysis of MG data. Each point indicates each analyzed TF, horizontal axis indicates wPGSA t score, and vertical axis indicates log_10_ P.Val. (BH adjusted). Colored points indicate TFs with padj ≤ 10E-5 and |tscore| ≥4. (E) Venn diagram indicating numbers of TFs identified from MG and AST data with significant enrichment in wPGSA.

Expression profiles of MG and AST showed that there was more prominent perturbation of gene expression in MG than in AST (Table S1-2), most likely due to higher expression of LPS recognizing toll-like receptors in MG[33, 34]. Differential expression analysis identified 48 DEGS (upreg: 3, downreg:45) that were padj ≤ 0.05, |Fold Change| ≥2 in MG as a result of *Kpna4* deficiency (Table S3). Notably, a larger number of downregulated genes compared to upregulated genes were identified, suggesting general repression of gene expression in both MG (Fig 4B; Table S1,S3) and AST (Fig S9A; Table S2,S4). As depletion of nuclear transport factors is likely to disrupt nuclear localization of specific transcription factors (TFs) or alter chromatin states[21], we sought to identify TFs and histone modifications upstream of altered genes by gene set enrichment analysis in Enrichr.[39–41] By individually analyzing upregulated and downregulated gene sets of p ≤ 0.05, |Fold Change| ≥2 (Table S5-14), we found significant enrichment only in the MG downregulated gene set (792 genes) (Fig 4C), where significant enrichment of Polycomb repressor complex 2 (PRC2) component TFs (SUZ12, EZH2 and JARID2) (Fig 4C left, Table S8), as well as repressive histone modifications introduced by PRC2 (Fig 4C right, Table S10) was seen, suggesting aberrant PRC2 activation and global downregulation of PRC2 target genes. To further validate these results, we utilized another analysis method: weighted parametric gene set analysis (wPGSA)[42], which allows for the prediction of altered TFs directly from gene expression data, independent of gene sets. Application of wPGSA to MG expression data revealed prominent and significant enrichment of binding sites of PRC2 related TFs (EZH2, SUZ12, JARID2, EZH1, PHF19, RING2, etc.) in genes downregulated by *Kpna4* deficiency (t score ≤ 0), as well as suggesting a global downregulation of gene expression (i.e. the majority of TFs were t score ≤ 0) (Fig 4D, Table S15). Moreover, the majority of TFs enriched in MG, including PRC2 component TFs EZH2 and SUZ12, were common between our MG (Table S15) and AST (Table S16) wPGSA results (Fig 4E, Fig S9 B), suggesting that altered PRC2 activity may underlie the increased inflammatory responses observed in KPNA4 cells in this study.

Finally, we performed cross-validation for our results with a previously reported dataset from *Kpna4* deficient neural cells (Dorsal Root Ganglion, Marvaldi et al[30]), and found similar enrichment of PRC2 component TFs in genes downregulated by *Kpna4* deficiency following tissue damage (Day7; Fig S9 E-F, Table S18). Such results support that *Kpna4* deficiency increases PRC2 activity and suppresses global gene expression.

## DISCUSSION

In this study, we demonstrated for the first time that a importin α4 (*Kpna4*) deficient mouse line exhibits increased anxiety-related behaviors, decreased social interaction, decreased avoidance learning, and decreased prepulse inhibition. As the previous study associating *KPNA4* to schizophrenia in postmortem samples had suggested contributions of *KPNA4* deficiency in the downregulation of NF-κB pathways in patients[6], we assessed the expression of NF-κB genes as well as proinflammatory cytokines downstream of the NF-κB pathway downstream in KO mice. Contrary to initial predictions based on the report[6], we found that *Kpna4* deficiency did not decrease NF-κB nuclear localization, but instead causes an increase, along with increased proinflammatory responses in KO tissues and cultured cells. Moreover, such proinflammatory increase was rescued by addition of not only full length, but also transport deficient (ΔIBB) mutants. We have previously demonstrated that this ΔIBB mutant of importin α can migrate to the nucleus by independently of importin β1 and Ran[43]. In addition, our immunohistochemical analysis demonstrated that importin α4 was observed in the nucleus of mouse brain tissue, suggesting that disruption of chromatin regulatory functions, not nuclear transport disfunction, from importin α4 deficiency induces increased proinflammatory responses in correlation to abnormal behavior in mice. Finally, we examined gene expression in sorted adult glial cells from KO mice, finding evidence supporting increased PRC2 activity which may underlie the perturbations in gene expression caused by *Kpna4* deficiency.

In this study, we found altered behavioral characteristics in KO mice in anxiety-like behavior (OFT, EPM), social interaction, passive avoidance (IA), and PPI. This is in line with the initial report in Roussos and collegues[6] that *KPNA4* deficiency associates to higher risks of schizophrenia and decreased PPI. Two previous studies[8, 30] have performed behavioral testing on KO mice. In particular, Panayotis, et. al.[8] has examined three behaviors (locomotion, anxiety-related behavior, and acoustic startle response), and have found alterations in home cage activity (higher and lower in KO than in WT in dark and light phases, respectively), no alterations in anxiety-like behavior, and increased startle response. Moreover, Malvaldi et al.[30] has found decreased pain responses in KO mice. The discrepancy between our results and the past studies may be due to differences in experimental protocol, where the past studies have performed behavioral analysis during the “dark” phase of the light cycle, as well as differences in background substrains of the KO mice used (C57BL/6JJcl vs C57BL/6OlaHsd). Notably, as KO mice has been reported to exhibit decreased pain responses, the decreased latency to enter the shock paired chamber observed in our IA experiments may be due to attenuated pain response instead of deficits in avoidance learning.

From examination of schizophrenia postmortem patients, Roussos et al[6] found NF-κB downregulation and significant association of KPNA4 genetic elements in the patient group, leading to the prediction that *KPNA4* deficiency results in the depletion of NF-κB factors from the nucleus in schizophrenia patients, and downregulation of NF-κB downstream genes including proinflammatory cytokines. This is contrastive to the inflammation hypothesis of schizophrenia, arising from observations that patients show immunoinflammatory upregulation, especially during acute psychiatric exacerbation[44–47]. In respect to this, it was speculated that the downregulated NF-κB signaling in postmortem brains (chronic patients) may have resulted from compensatory NF-κB pathway alteration as a result of initial activation in acute patient[6]. In our examination of NF-κB and proinflammatory cytokine expression in *Kpna4* deficient mice of a relatively young age (8-10 weeks), we observed an increase in *RelA* mRNA expression, as well as repeated evidence of an increase in proinflammatory cytokines when presented with immunological stimulation in cells collected from embryonic or relatively young (10 weeks) mice. This observation supports possibilities that *KPNA4* deficiency may cause increased NF-κB signaling in younger patients in the acute phase of schizophrenia, which is in line with the inflammation hypothesis. Examination of NF-κB signaling and neuroinflammation in aged *Kpna4* deficient mice, as well as insight into the relationship between KPNA4 and cellular senescence may provide more insight on these age-related discrepancies.

Considering that *Kpna4* deficiency alone did not result in exclusion of p65 from the nucleus in mice, contribution from redundant nuclear import pathways may sustain the import of nuclear proteins essential for central cellular functions such as inflammation. There is extensive evidence demonstrating redundancy in both classical[25, 26, 48, 49] and non-classical[49] nuclear import of NF-κB proteins, and depletion of *Kpna4* may not be sufficient to hinder the import of a specific nuclear protein. Near-complete exclusion of essential TFs from the nucleus seems to be lethal, as shown in our report where mice deficient in both *Kpna4* and *Kpna3* (known as major transporters of NF-κB[25]) were not viable[21]. Moreover, as transfection with a nuclear transport deficient mutant (ΔIBB) was sufficient for reversing the proinflammatory increase in this study, such increases seen in this study may be a result of the direct gene regulatory functions, rather than the nuclear transport functions, of importin αs.

In our analysis of gene expression perturbations in *Kpna4* deficient MG and Ast cells, we found prominent enrichment of binding sites for PRC2 components (EZH2, SUZ12, JARID2, etc.) in genes downregulated in KO. PRC2 is a chromatin modifying complex mainly known for repressive H3K27 methylation and has a plethora of roles in cellular functions such as neural differentiation, immunoinflammatory regulation, and tumor regulation[50], and its enrichment in downregulated genes suggest that *Kpna4* deficiency increases PRC2 activity. KPNA4 has been identified as a binding partner for PRC2 component EZH2[51], and its cellular depletion may alter EZH2 functionality to increase repressive activity of PRC2. Moreover, we revealed that rescue of the ΔIBB mutant was sufficient in reversing increases in proinflammatory responses, which implies that the immunosuppressive effects of importin α4 are mediated through direct gene regulation function, rather than its transport functions. Furthermore, decreased active states of chromatin were implied by proteomics analysis, which may be explained by increased PRC2 activity causing aberrant silencing of widespread target loci[21]. In regard to increased immunoinflammatory activation in KO glial cells, we observed enrichment of TFs responsible for immunoinflammatory activation (FLI1, ELF1, ASH2L, ETV6) upstream of genes upregulated in KO cells. Increased PRC2 activity has been found to alter microglial polarization towards a proinflammatory (M1-like) status and induces upregulation of proinflammatory cytokines[52], which has been observed in our KO cells. The precise molecular interactions that mediate PRC2 hyperactivity in *Kpna4* deficient cells and the implications in regulation of behavior is still unknown and calls for further detailed examination; however, there is accumulating evidence that increased immunoinflammatory activity[44, 53–55], as well as repression of gene expression (decreased gene expression[56], increase in histone deacetylase and/or repressive histone modifications[57–59]) are involved in the pathology of schizophrenia and/or schizophrenia-associated behaviors, and KO mice may provide a useful tool to understand the molecular basis of behavioral dysregulation triggered by neuroinflammation.

Significant enrichment of AP1 family factors upstream of genes downregulated by *Kpna4* deficiency, as well as decreased nuclear localization of c-fos in peripheral sensory neurons caused by *Kpna4* deficiency has been demonstrated previously.[30]; however, in our experiments we were not able to observe enrichment of c-fos or other AP1 factors in our analysis of expression patterns from FACS sorted adult glial cells. This may suggest that specific regulatory roles of importin α4 exist between different neural cell types, which are defined by varying expression patterns of different nuclear transport factors and cargo. Further studies are required for understanding the correlation between the expression and function of importin α4 in different cells and tissues.

The results from this study emphasize the roles of importin α4 in behavioral and neuroinflammatory regulation in relation to psychiatric disorders. Furthermore, as two other importin α subtypes (*Kpna1*[1, 2] and *Kpna3*[3–5]) have been associated to psychiatric disorders, further understanding and comparison of subtype specific cargo binding capabilities and non-transport functions may be important in uncovering the molecular pathology of psychiatric disorders. Additionally, as exportin 7, an importinβ family nuclear transport factor, is coded in one of the top schizophrenia-associated loci[60], further insight into the relationship between nucleocytoplasmic transport and behavioral regulation is necessary to undercover the functions of nuclear import factors in the pathology of psychiatric disorders.

## MATERIALS AND METHODS

### Animals

Heterozygous (Het), and homozygous Importin α4 (*Kpna4*) knockout (KO), as well as wild type (WT) mice on a C57BL6/JJcl (CLEA Japan Inc., Tokyo, Japan) background were generated by mating male and female Kpna4 Het mice[21]. All animal experiments complied with institutional guidelines by the Institutional Safety Committee on Recombinant DNA Experiments (No. 04219 and 04884) at Osaka University, (No. 110083) and (No. DNA-420) at NIBIOHN, Animal Experimental Committee of the Institute for Protein Research at Osaka University (No. 29-02-1 and R04-01-1), the Animal Care and Use Committee of Kyoto University (No. MedKyo17071), and animal research committees of NIBIOHN (No. DS26-34).

### Behavioral Tests

The behavioral tests were administered to two different cohorts of mice, with cohort 1 being administered the following tests open field test (OFT), elevated plus maze (EPM), Y-Maze, social interaction test, inhibitory avoidance (IA), administered in the above order. Cohort 2 was administered the prepulse inhibition (PPI) test and used for tissue collection (RNA extraction). Behavioral tests were administered following previously reported general procedures with minor modifications.[9, 10, 31, 32, 61]

Detailed protocols on behavioral testing, along with dissection, qRT-PCR, SDS-PAGE and immunoblotting, cell culture, and RNAseq analysis are described in the supplementary materials. Data files from our RNAseq experiments have been submitted to Gene Expression Omnibus (GEO) as GSE264180.

## ACKNOWLEDGEMENTS

We thank Toshie Deki, Elizabeth Richards, Noriko Otani, and Taichi Itou for technical assistance.

## CONTRIBUTIONS

MOka, TH, and YM contributed to the conception of the study. KS, MMorita, YA, MOka, TH, and YM designed experiments. KS, MMorita, MOtani, RO, KL, and YM performed data acquisition and analysis. KS, YA, TH, and YM interpreted data. KS and MMorita performed behavioral analysis. KS and YA performed RNAseq and wPGSA analysis. KS and MMatsumoto performed MACS and FACS isolation of glial cells. EK and AS contributed to multiplex Immunoarray analysis. TM, AS, KL, MY, YY, MOka, TH, and YM supervised the study. KS, MOka TH, and YM wrote the manuscript. All authors contributed to manuscript revision and read and approved the submitted version.

## FUNDING

This work was supported in part by the JSPS KAKENHI Grants (JP15K07068, JP20K06455, and JP22KK0111 to YM, 16H04789 to YY and MOka, JP22H02944 and JP23K18163 to TH, and 17H03679, 20H03444, 23K27373 to MOka), AMED Grants (JP21wm0425010 and 21gm1510006 to TH), JST SPRING (JPMJSP2138 to KS and YA), AMED BINDS Research Support Project for Life Science and Drug Discovery (23ama121052 and JP23ama121054), Salt Science Research Foundation Grants (2229 to TH), the Collaborative Research Program of the Institute for Protein Research (ICR-24-03), and Institute for Protein Research Promotion Program for Frontier Protein Research (to KS).

## COMPETING INTERESTS

The authors declare no competing interests.

## Notes

### Competing Interest Statement

The authors have declared no competing interest.

## References

1. Girard SL, Gauthier J, Noreau A, Xiong L, Zhou S, Jouan L, et al. Increased exonic de novo mutation rate in individuals with schizophrenia. Nat Genet. 2011;43:860–863.

2. Jouan L, Girard SL, Dobrzeniecka S, Ambalavanan A, Krebs M-O, Joober R, et al. Investigation of rare variants in LRP1, KPNA1, ALS2CL and ZNF480 genes in schizophrenia patients reflects genetic heterogeneity of the disease. Behav Brain Funct. 2013;9:9.

3. Wei J, Hemmings GP. The KPNA3 gene may be a susceptibility candidate for schizophrenia. Neurosci Res. 2005;52:342–346.

4. Zhang H, Ju G, Wei J, Hu Y, Liu L, Xu Q, et al. A combined effect of the KPNA3 and KPNB3 genes on susceptibility to schizophrenia. Neurosci Lett. 2006;402:173–175.

5. Morris CP, Baune BT, Domschke K, Arolt V, Swagell CD, Hughes IP, et al. KPNA3 variation is associated with schizophrenia, major depression, opiate dependence and alcohol dependence. Dis Markers. 2012;33:163–170.

6. Roussos P, Katsel P, Davis KL, Giakoumaki SG, Siever LJ, Bitsios P, et al. Convergent Findings for Abnormalities of the NF-κB Signaling Pathway in Schizophrenia. Neuropsychopharmacol. 2013;38:533.

7. Hosokawa K, Nishi M, Sakamoto H, Tanaka Y, Kawata M. Regional distribution of importin subtype mRNA expression in the nervous system: Study of early postnatal and adult mouse. Neuroscience. 2008:864–877.

8. Panayotis N, Sheinin A, Dagan SY, Tsoory MM, Rother F, Vadhvani M, et al. Importin α5 Regulates Anxiety through MeCP2 and Sphingosine Kinase 1. Cell Reports. 2018;25:3169–3179.e7.

9. Sakurai K, Itou T, Morita M, Kasahara E, Moriyama T, Macpherson T, et al. Effects of Importin α1/KPNA1 deletion and adolescent social isolation stress on psychiatric disorder-associated behaviors in mice. Plos One. 2021;16:e0258364.

10. Nomiya H, Sakurai K, Miyamoto Y, Oka M, Yoneda Y, Hikida T, et al. A Kpna1-deficient psychotropic drug-induced schizophrenia model mouse for studying gene–environment interactions. Sci Rep. 2024;14:3376.

11. Aomine Y, Sakurai K, Macpherson T, Ozawa T, Miyamoto Y, Yoneda Y, et al. Importin α3 (KPNA3) Deficiency Augments Effortful Reward-Seeking Behavior in Mice. Front Neurosci-Switz. 2022;16:905991.

12. Seki T, Tada S, Katada T, Enomoto T. Cloning of a cDNA Encoding a Novel Importin-α Homologue, Qip1: Discrimination of Qip1 and Rch1 from hSrp1 by Their Ability to Interact with DNA Helicase Q1/RecQL. Biochem Biophys Res Commun. 1997;234:48–53.

13. Wälde S, Kehlenbach RH. The Part and the Whole: functions of nucleoporins in nucleocytoplasmic transport. Trends Cell Biol. 2010;20:461–469.

14. D’Angelo MA, Hetzer MW. Structure, dynamics and function of nuclear pore complexes. Trends Cell Biol. 2008;18:456–466.

15. Goldfarb DS, Corbett AH, Mason DA, Harreman MT, Adam SA. Importin alpha: a multipurpose nuclear-transport receptor. Trends Cell Biol. 2004;14:505–514.

16. Pumroy RA, Cingolani G. Diversification of importin-α isoforms in cellular trafficking and disease states. Biochem J. 2015;466:13–28.

17. Jans DA, Xiao C, Lam MHC. Nuclear targeting signal recognition: a key control point in nuclear transport? Bioessays. 2000;22:532–544.

18. Sekimoto T, Yoneda Y. Intrinsic and extrinsic negative regulators of nuclear protein transport processes. Genes Cells. 2012;17:525–535.

19. Yasuda Y, Miyamoto Y, Yamashiro T, Asally M, Masui A, Wong C, et al. Nuclear retention of importin α coordinates cell fate through changes in gene expression. Embo J. 2012;31:83–94.

20. Miyamoto Y, Yamada K, Yoneda Y. Importin α: a key molecule in nuclear transport and non-transport functions. J Biochem. 2016;160:69–75.

21. Miyamoto Y, Sasaki M, Miyata H, Monobe Y, Nagai M, Otani M, et al. Genetic loss of importin α4 causes abnormal sperm morphology and impacts on male fertility in mouse. Faseb J. 2020. 2020. 10.1096/fj.202000768rr.

22. Oka M, Yoneda Y. Importin α: functions as a nuclear transport factor and beyond. Proc Jpn Acad Ser B. 2018;94:259–274.

23. Panayotis N, Karpova A, Kreutz MR, Fainzilber M. Macromolecular transport in synapse to nucleus communication. Trends Neurosci. 2015;38:108–116.

24. Baker SA, Lombardi LM, Zoghbi HY. Karyopherin α 3 and karyopherin α 4 proteins mediate the nuclear import of methyl-CpG binding protein 2. J Biological Chem. 2015;290:22485–22493.

25. Fagerlund R, Kinnunen L, Köhler M, Julkunen I, Melén K. NF-κB Is Transported into the Nucleus by Importin α3 and Importin α4. J Biol Chem. 2005;280:15942–15951.

26. Fagerlund R, Melén K, Cao X, Julkunen I. NF-κB p52, RelB and c-Rel are transported into the nucleus via a subset of importin α molecules. Cell Signal. 2008;20:1442–1451.

27. Agrawal T, Gupta GK, Agrawal DK. Effect Of Cytokines On Expression Of Importin Alpha3 (KPNA4) In Human Bronchial Smooth Muscle Cells. American Thoracic Society 2011 International Conference, May 13-18, 2011 • Denver Colorado. 2011:A2580–A2580.

28. Theiss AL, Jenkins AK, Okoro NI, Klapproth J-MA, Merlin D, Sitaraman SV. Prohibitin Inhibits Tumor Necrosis Factor alpha–induced Nuclear Factor-kappa B Nuclear Translocation via the Novel Mechanism of Decreasing Importin α3 Expression. Mol Biol Cell. 2009;20:4412–4423.

29. Thiele S, Stanelle-Bertram S, Beck S, Kouassi NM, Zickler M, Müller M, et al. Cellular Importin-α3 Expression Dynamics in the Lung Regulate Antiviral Response Pathways against Influenza A Virus Infection. Cell Reports. 2020;31:107549.

30. Marvaldi L, Panayotis N, Alber S, Dagan SY, Okladnikov N, Koppel I, et al. Importin α3 regulates chronic pain pathways in peripheral sensory neurons. Sci New York N Y. 2020;369:842–846.

31. Ikeda K, Sato A, Mizuguchi M. Social interaction test: a sensitive method for examining autism-related behavioral deficits. Protoc Exch. 2013. 2013. 10.1038/protex.2013.046.

32. Hikida T, Kimura K, Wada N, Funabiki K, Nakanishi S. Distinct Roles of Synaptic Transmission in Direct and Indirect Striatal Pathways to Reward and Aversive Behavior. Neuron. 2010;66:896–907.

33. Uhlén M, Björling E, Agaton C, Szigyarto CA-K, Amini B, Andersen E, et al. A Human Protein Atlas for Normal and Cancer Tissues Based on Antibody Proteomics. Mol Cell Proteom. 2005;4:1920–1932.

34. Sjöstedt E, Zhong W, Fagerberg L, Karlsson M, Mitsios N, Adori C, et al. An atlas of the protein-coding genes in the human, pig, and mouse brain. Science. 2020;367.

35. Talcott B, Moore MS. The Nuclear Import of RCC1 Requires a Specific Nuclear Localization Sequence Receptor, Karyopherin α3/Qip*. J Biol Chem. 2000;275:10099–10104.

36. Köhler M, Speck C, Christiansen M, Bischoff FR, Prehn S, Haller H, et al. Evidence for Distinct Substrate Specificities of Importin α Family Members in Nuclear Protein Import. Mol Cell Biol. 1999;19:7782–7791.

37. Zhu J, Cynader MS, Jia W. TDP-43 Inhibits NF-κB Activity by Blocking p65 Nuclear Translocation. PLoS ONE. 2015;10:e0142296.

38. Liu L, McBride KM, Reich NC. STAT3 nuclear import is independent of tyrosine phosphorylation and mediated by importin-α3. P Natl Acad Sci Usa. 2005;102:8150–8155.

39. Kuleshov MV, Jones MR, Rouillard AD, Fernandez NF, Duan Q, Wang Z, et al. Enrichr: a comprehensive gene set enrichment analysis web server 2016 update. Nucleic Acids Res. 2016;44:W90–W97.

40. Chen EY, Tan CM, Kou Y, Duan Q, Wang Z, Meirelles GV, et al. Enrichr: interactive and collaborative HTML5 gene list enrichment analysis tool. BMC Bioinform. 2013;14:128.

41. Xie Z, Bailey A, Kuleshov MV, Clarke DJB, Evangelista JE, Jenkins SL, et al. Gene Set Knowledge Discovery with Enrichr. Curr Protoc. 2021;1:e90.

42. Kawakami E, Nakaoka S, Ohta T, Kitano H. Weighted enrichment method for prediction of transcription regulators from transcriptome and global chromatin immunoprecipitation data. Nucleic Acids Res. 2016;44:5010–5021.

43. Miyamoto Y, Hieda M, Harreman MT, Fukumoto M, Saiwaki T, Hodel AE, et al. Importin alpha can migrate into the nucleus in an importin beta- and Ran-independent manner. Embo J. 2002;21:5833–5842.

44. Miller BJ, Goldsmith DR. Evaluating the Hypothesis That Schizophrenia Is an Inflammatory Disorder. FOCUS. 2020;18:391–401.

45. Song X-Q, Lv L-X, Li W-Q, Hao Y-H, Zhao J-P. The Interaction of Nuclear Factor-Kappa B and Cytokines Is Associated with Schizophrenia. Biol Psychiat. 2009;65:481–488.

46. Hashimoto R, Ohi K, Yasuda Y, Fukumoto M, Yamamori H, Takahashi H, et al. Variants of the RELA gene are associated with schizophrenia and their startle responses. Neuropsychopharmacol Official Publ Am Coll Neuropsychopharmacol. 2011;36:1921–1931.

47. Frommberger UH, Bauer J, Haselbauer P, Fräulin A, Riemann D, Berger M. Interleukin-6-(IL-6) plasma levels in depression and schizophrenia: comparison between the acute state and after remission. Eur Arch Psy Clin N. 1997;247:228–233.

48. Tao R, Xu X, Sun C, Wang Y, Wang S, Liu Z, et al. KPNA2 interacts with P65 to modulate catabolic events in osteoarthritis. Exp Mol Pathol. 2015;99:245–252.

49. Liang P, Zhang H, Wang G, Li S, Cong S, Luo Y, et al. KPNB1, XPO7 and IPO8 Mediate the Translocation ofNF-κB/p65 into the Nucleus. Traffic. 2013;14:1132–1143.

50. Liu X, Liu X. PRC2, Chromatin Regulation, and Human Disease: Insights From Molecular Structure and Function. Front Oncol. 2022;12:894585.

51. Oliviero G, Brien GL, Waston A, Streubel G, Jerman E, Andrews D, et al. Dynamic Protein Interactions of the Polycomb Repressive Complex 2 during Differentiation of Pluripotent Cells. Mol Cell Proteomics. 2016;15:3450–3460.

52. Yang X, Zhang Y, Chen Y, He X, Qian Y, Xu S, et al. LncRNA HOXA-AS2 regulates microglial polarization via recruitment of PRC2 and epigenetic modification of PGC-1α expression. J Neuroinflammation. 2021;18:197.

53. Murphy CE, Walker AK, Weickert CS. Neuroinflammation in schizophrenia: the role of nuclear factor kappa B. Transl Psychiat. 2021;11:528.

54. Williams JA, Burgess S, Suckling J, Lalousis PA, Batool F, Griffiths SL, et al. Inflammation and Brain Structure in Schizophrenia and Other Neuropsychiatric Disorders. JAMA Psychiatry. 2022;79:498–507.

55. Fond G, Lançon C, Korchia T, Auquier P, Boyer L. The Role of Inflammation in the Treatment of Schizophrenia. Front Psychiatry. 2020;11:160.

56. Clifton NE, Hannon E, Harwood JC, Florio AD, Thomas KL, Holmans PA, et al. Dynamic expression of genes associated with schizophrenia and bipolar disorder across development. Transl Psychiatry. 2019;9:74.

57. Lang B, Alrahbeni TMA, Clair DS, Blackwood DH, Consortium IS, McCaig CD, et al. HDAC9 is implicated in schizophrenia and expressed specifically in post-mitotic neurons but not in adult neural stem cells. Am J Stem Cells. 2011;1:31–41.

58. Bahari-Javan S, Varbanov H, Halder R, Benito E, Kaurani L, Burkhardt S, et al. HDAC1 links early life stress to schizophrenia-like phenotypes. Proc Natl Acad Sci. 2017;114:E4686–E4694.

59. Ibi D, Revenga M de la F, Kezunovic N, Muguruza C, Saunders JM, Gaitonde SA, et al. Antipsychotic-induced Hdac2 transcription via NF-κB leads to synaptic and cognitive side effects. Nat Neurosci. 2017;20:1247–1259.

60. Singh T, Poterba T, Curtis D, Akil H, Eissa MA, Barchas JD, et al. Rare coding variants in ten genes confer substantial risk for schizophrenia. Nature. 2022:1–9.

61. Li S, Sakurai K, Ohgidani M, Kato TA, Hikida T. Ameliorative effects of Fingolimod (FTY720) on microglial activation and psychosis-related behavior in short term cuprizone exposed mice. Mol Brain. 2023;16:59.

