## Supplemental File for "Importin α4 deficiency induces psychiatric disorder-related behavioral deficits and neuroinflammation in mice"

*Corresponding Authors

#### Contents:


1. **Supplemental Figures**
2. **Detailed Materials and Methods**
3. **References for Supplemental Material**


(Statistical Analysis and Supplemental Tables are available as separate files.)

### 1. Supplemental Figures

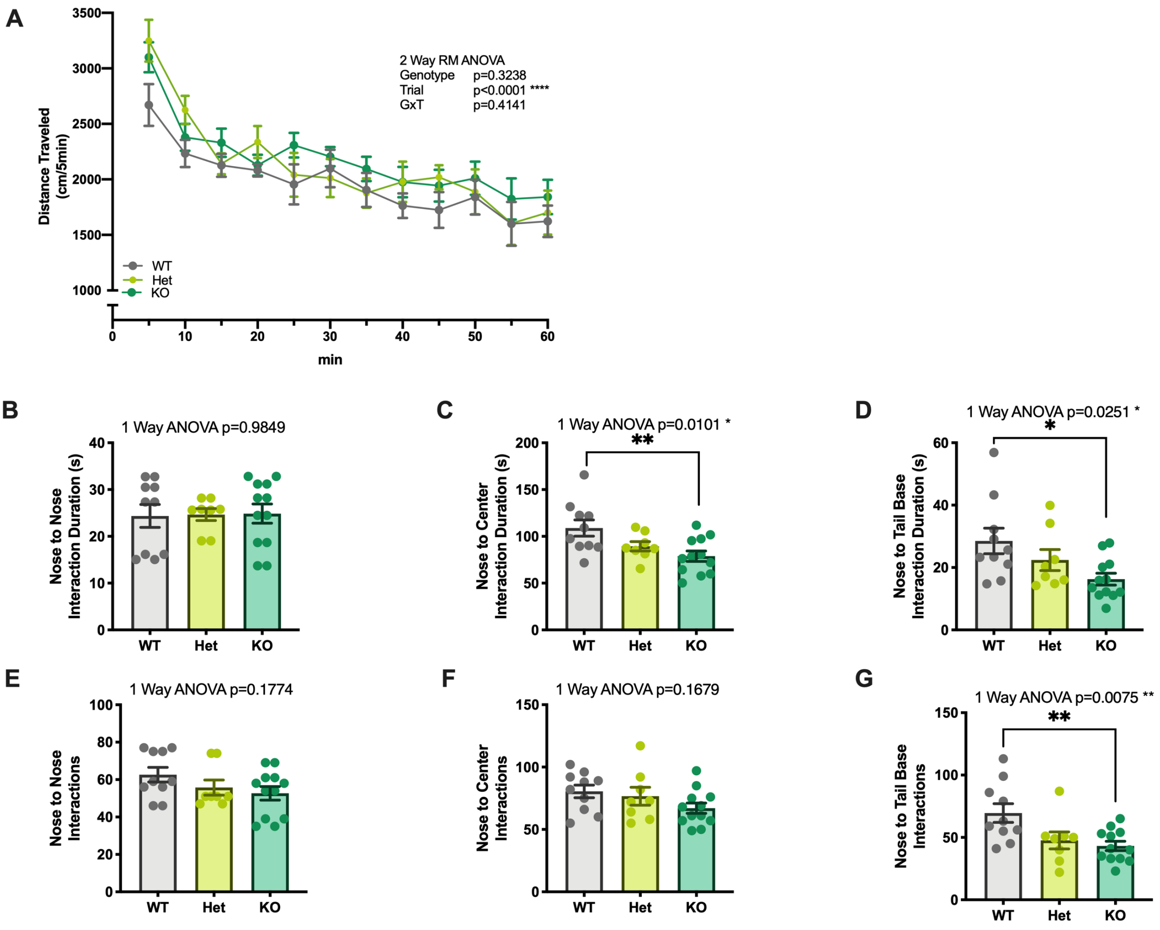


**Fig S1 Additional Measures from Behavioral Battery**

(A) Distance travelled per 5 minutes in the OFT. (B) Duration of Nose to Nose interactions in the SIT. (C) Duration of Nose to Center interactions in the SIT. (C) Duration of Nose to Tail Base interactions in the SIT. (E) Number of Nose to Nose interactions in the SIT. (F) Number of Nose to Center interactions in the SIT. (G) Number of Nose to Tail Base interactions in the SIT. (A) Each data point represents mean distance (per 5min), error bars represent SEM. (B-G) Each bar represents mean, error bars represent SEM. Post-hoc Tukey Test: **p<0.01, *p<0.05.

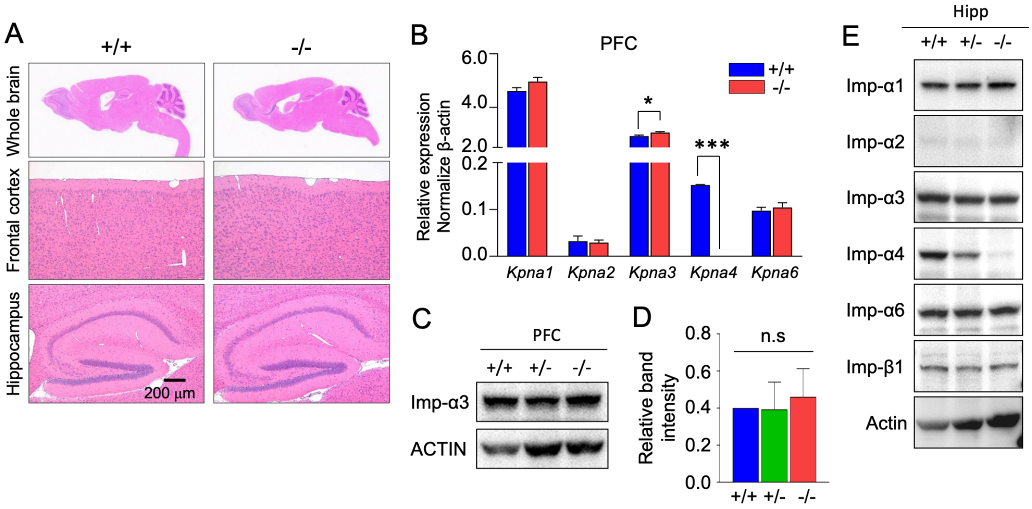


#### Fig S2 Morphology and expression levels of other Importin α subtypes in KO brains

(A) Hematoxylin and eosin (HE)-stained sagittal sections of paraffin-embedded mouse brains from 50-day-old WT and KO mice. Images are from the whole brain, prefrontal cortex and hippocampus. Scale bars: 200 μm. (B) Transcript levels of importin α subtypes (*Kpna1, Kpna2, Kpna3, Kpna4,* and *Kpna6*) were measured in the KO PFCs. The β-actin gene was used as an internal control. Data are presented as fold-change compared to WT PFCs (n=3 each, mean ± SEM). Statistical significance was determined using two-way ANOVA and Post-hoc Sidak Test, where $^{*}P<0.05$, $^{***}P<0.001$. (C-D) Western blot of KPNA3 on KO PFC. Actin was used as an internal control. (E) Western blot of importin α subtypes (Importin αs: Imp-α1 (KPNA1), Imp-α2 (KPNA2), Impα-3 (KPNA3), Imp-α4 (KPNA4), and Imp-α6 (KPNA6)) and importin β1 (Imp-β1 (KPNB1) in the KO hippocampus. Actin was used as an internal control.

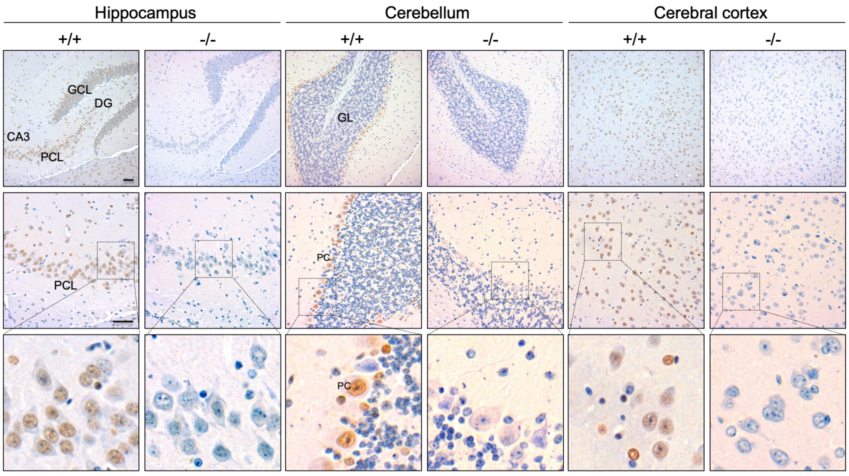


#### Fig S3 Staining of Importin α4 in WT and KO brains

Staining of Importin α4 in the Hippocampus, Cerebellum and prefrontal cortex of WT($^{+/+}$) and KO brains. DG: Dentate Gyrus, GCL: Granule Cell layer, CA: Cornu Ammonis, PCL: Pyramidal cell layer, PC: Purkinje cell, GL: Granule layer. Scale bars = 50μm.

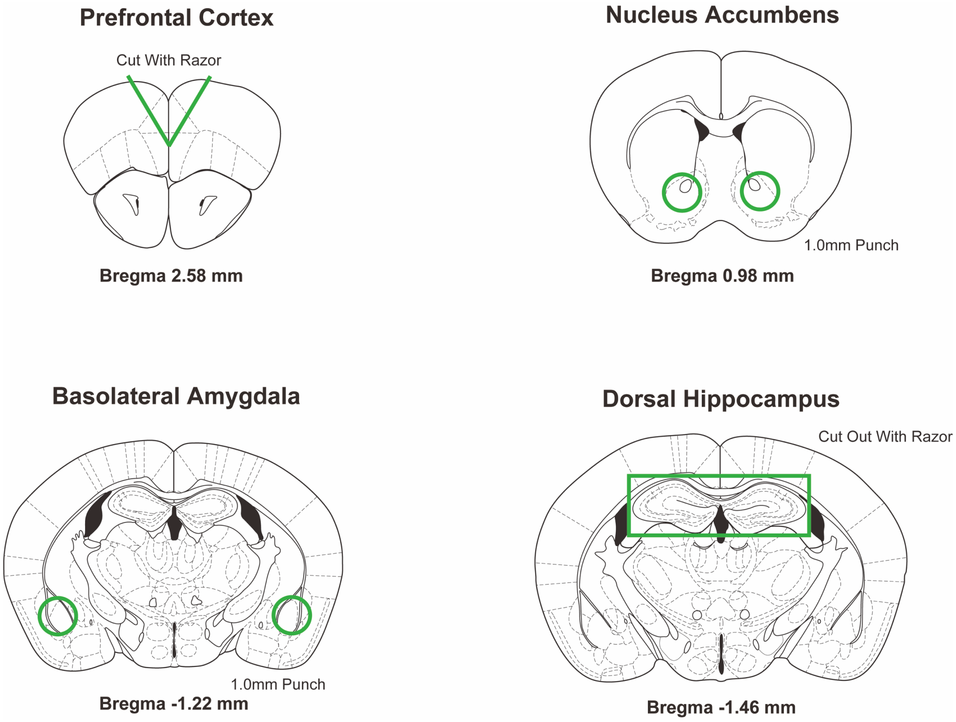


**Fig S4 Coordinates for Brain Tissue Samples**

Coordinates used for brain tissue sampling.

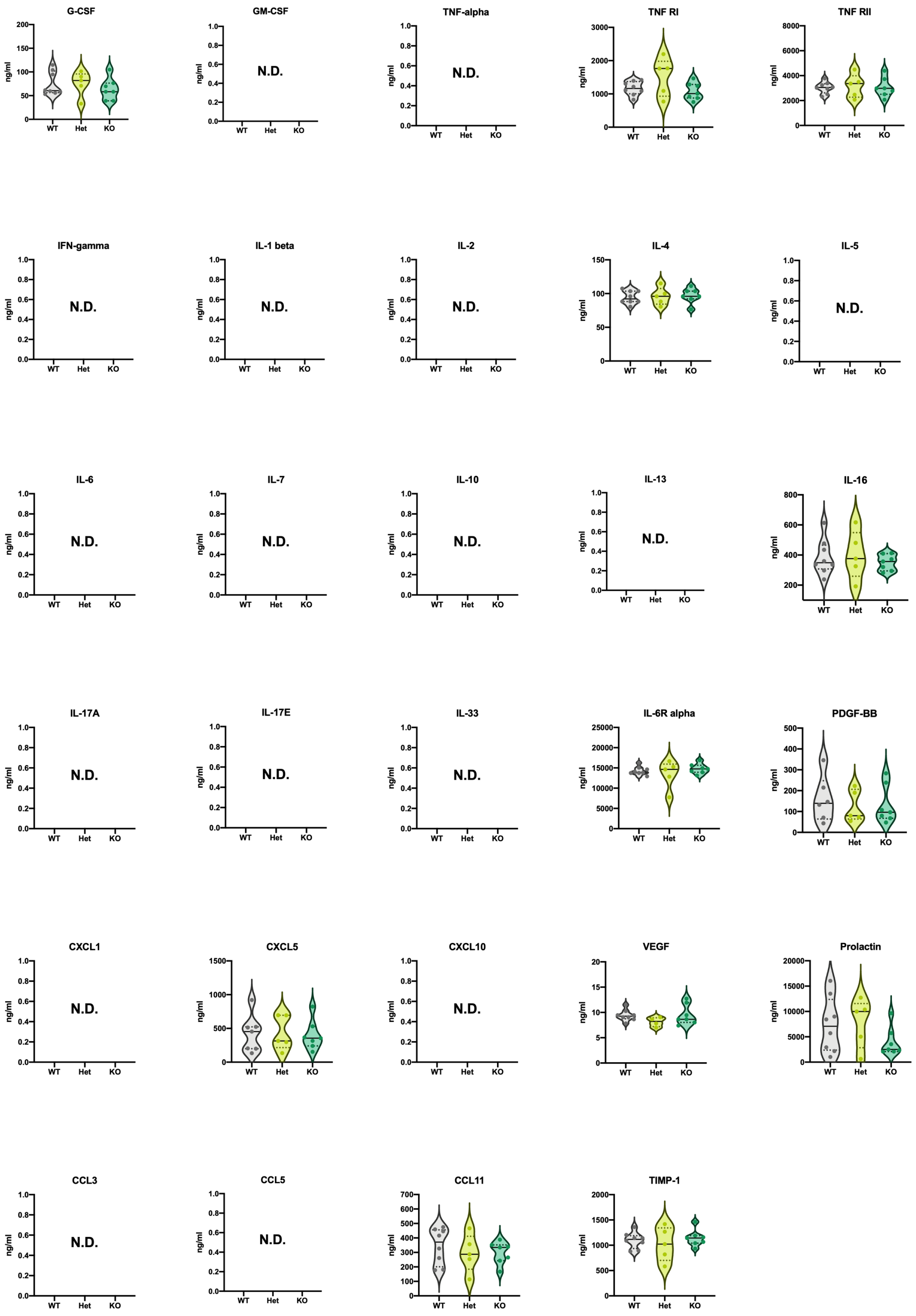


#### Fig S5 Analysis of Plasma in Multiplex Immunoarray Panel

Assessment of plasma protein levels in WT (n=8), Het (n=5), and KO (n=7) mice using a multiplex bead suspension Immunoarray panel to assess cytokines, chemokines, and other signaling proteins. Protein concentration was measured in ng/ml of plasma, with violin plots indicating median (solid line), and quartiles (dotted line). Data points which were below the minimum limit of quantitation or determined as outliers in ROUT test (Q:1%; Graphpad Prism) are excluded from the plots, and N.D. indicates that all samples were below the minimum limit of quantitation.

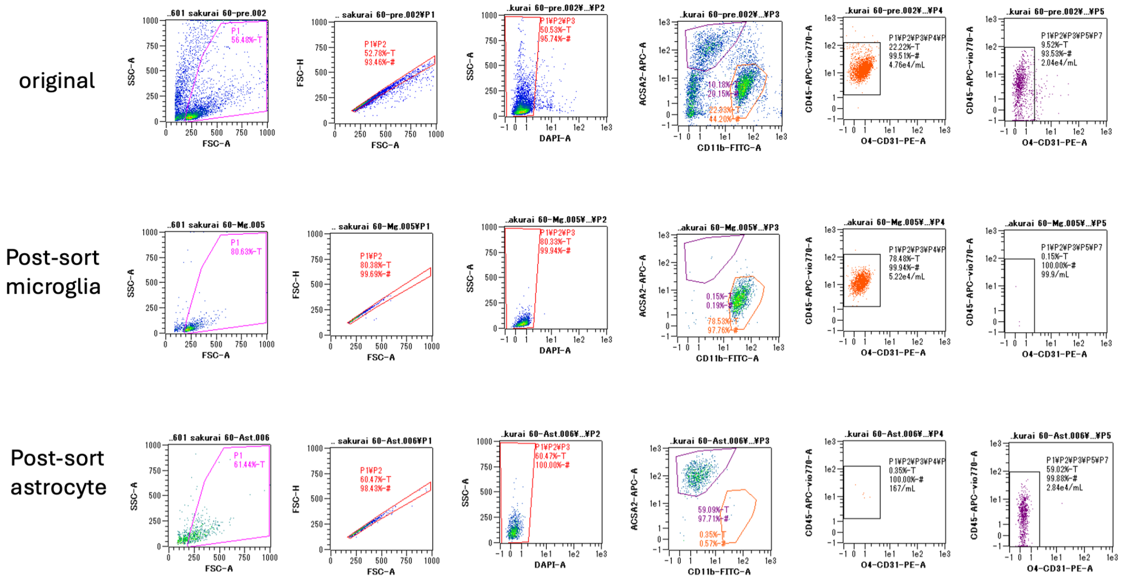


**Fig S6 FACS Sorting**

Representative sample of measurements for cell purity assessment using flow cytometry after FACS sorting of MG and AST.

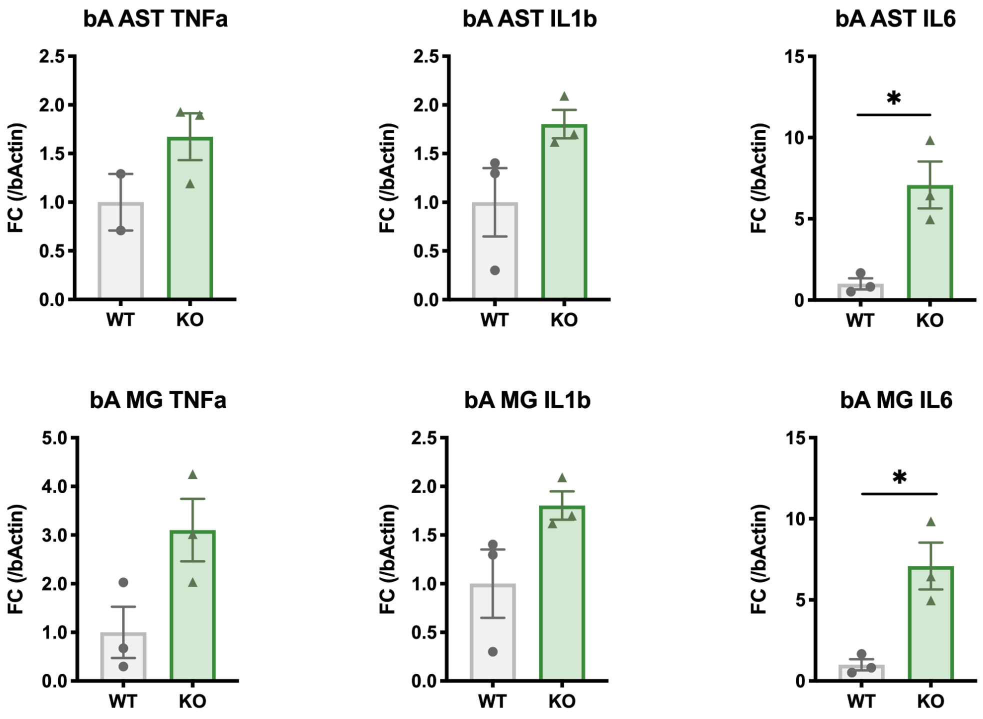


**Fig S7 Expression levels of TNF-a, IL-1b, and IL-6 in MACS isolated Astrocytes and Microglia from LPS injected Mice**

RT-qPCR analysis on expression levels of *Tnf-α*, *Il1-β*, and *Il-6* (/β-Actin) in Microglia and Astrocytes isolated from KO mice whole brain, injected with LPS following the same conditions as individuals used for RNAseq. Each bar represents mean, error bars represent SEM. Student's t-test *P<0.05.

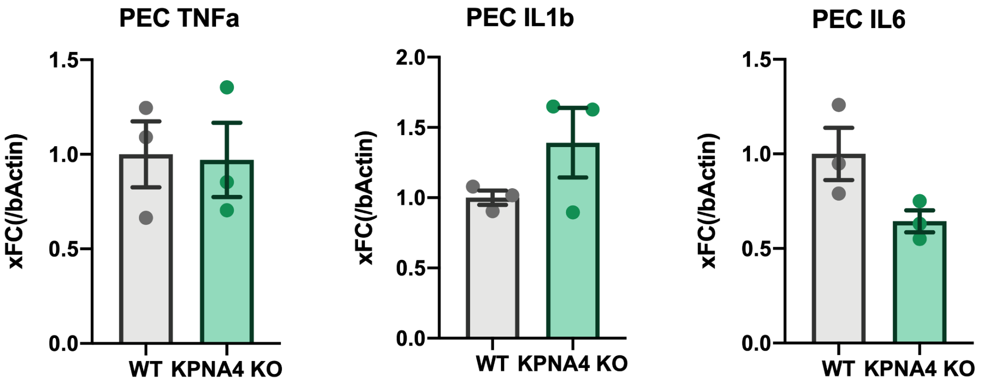


**Fig S8 Expression levels of TNF-a, IL-1b, and IL-6 in peritoneal macrophages collected from LPS injected Mice**

Expression levels of *Tnf-α*, *Il1-β*, and *Il-6* (/β-Actin) in peritoneal macrophages collected from KO mice injected with LPS used for RNAseq. Each bar represents mean, error bars represent SEM.

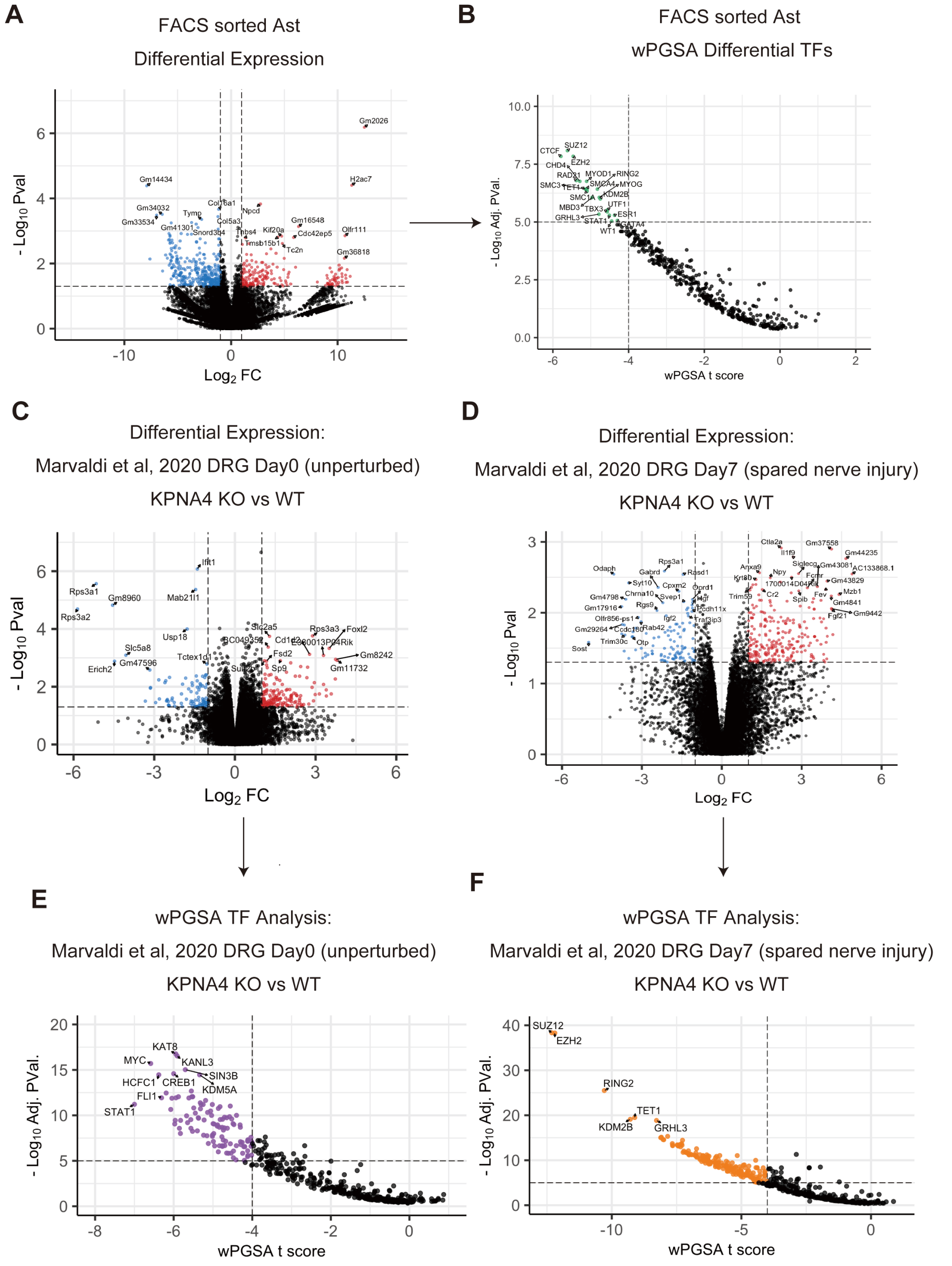


**Fig S9 RNAseq Analysis: AST and other Datasets**

(A) Volcano plot of genes with altered expression in KO ASTs in comparison to WT mice. Horizontal axis indicates log2 Fold Change, and vertical axis indicates -log10 p value. Colored points indicate genes with $p \leq 0.05$ and |Fold Change| ≥2. (B) Volcano plot of wPGSA analysis of AST data. Each point indicates each analyzed transcription factor, horizontal axis indicates wPGSA t score (positive and negative values indicate upregulation and downregulation, respectively, of genes with binding sites of indicated transcription factors), and vertical axis indicates log10 Adj.P.Val. (Benjamini-Hochberg adjusted). Colored points indicate TFs with padj ≤ 10E-5 and |tscore| ≥4. (C) Volcano plot of genes with altered expression in created from the dataset reported in Marvaldi et al[1], of KO Dorsal Root Ganglions in comparison to WT mice at day 0 (unperturbed). Horizontal axis indicates log2 Fold Change, and vertical axis indicates -log10 p value. Colored points indicate genes with p ≤ 0.05 and |Fold Change| ≥2. (D) Volcano plot of wPGSA analysis data from (C). Each point indicates each analyzed transcription factor, horizontal axis indicates wPGSA t score, and vertical axis indicates log10 Adj.P.Val. (BH adjusted). Colored points indicate TFs with padj ≤ 10E-5 and |tscore| ≥4. (E) Volcano plot of genes with altered expression in created from the dataset reported in Marvaldi et al[1], of KO Dorsal Root Ganglions in comparison to WT mice at day 7 after injury (spared nerve injury model). Horizontal axis indicates log2 Fold Change, and vertical axis indicates -log10 p value. Colored points indicate genes with p ≤ 0.05 and |Fold Change| ≥2. (F) Volcano plot of wPGSA analysis data from (E). Each point indicates each analyzed transcription factor, horizontal axis indicates wPGSA t score, and vertical axis indicates log10 Adj.P.Val. (BH adjusted). Colored points indicate TFs with padj ≤ 10E-5 and |tscore| ≥4.

#### 2. Detailed Materials and Methods

**Animals**

Heterozygous (Het), and homozygous Importin α4 (*Kpna4*) knockout (KO), as well as wild type (WT) mice on a C57BL6/JJcl (CLEA Japan Inc., Tokyo, Japan) background were generated at the Institute for Protein Research, Osaka University, by mating male and female *Kpna4* Het mice. The generation of *Kpna4* KO mice and genotyping methods have been previously reported[2]. Mice were housed on a 12-h light/dark cycle (Light: 0800–2000, Dark: 2000–0800) in a sound attenuated and air-conditioned room with temperatures maintained at 24 ± 2 °C. All mice were weaned and housed with their same-sex littermates (3–6 mice per cage) in standard cages (21 x 32 x 13 cm), with ad libitum access to standard mouse chow and drinking water. All animal experiments complied with institutional guidelines by the Institutional Safety Committee on Recombinant DNA Experiments (No. 04219 and 04884) at Osaka University, (No. 110083) and (No. DNA-420) at NIBIOHN, Animal Experimental Committee of the Institute for Protein Research at Osaka University (No. 29-02-1 and R04-01-1), the Animal Care and Use Committee of Kyoto University (No. MedKyo17071), and animal research committees of NIBIOHN (No. DS26-34).

**Behavioral Tests**

Behavioral tests were started after the mice reached 9 weeks of age. All tests were conducted during daytime (13:00–18:00) in the "light" phase light/dark cycle, in order to comply with institutional regulations. A total of 78 male mice (Cohort 1: WT 11, *Kpna4*$^{+/-}$ 8, and KO 12; Cohort 2: WT 16, Het 18, and KO 13) were subjected to the behavioral test battery (additional Statistical Analysis File). The behavioral tests were administered to two different cohorts of mice, with cohort 1 being administered the following tests open field test (OFT), elevated plus maze (EPM), Y-Maze, social interaction test, inhibitory avoidance (IA), administered in the above order. Cohort 2 was administered the prepulse inhibition (PPI) test and used for blood collection and/or RNA extraction experiments. All behavioral tests were administered with 2 to 5 days in between tests, at the Medical Innovation Center, Kyoto University following previously reported general procedures.[3–5] Experimenters performing experiments and manual analysis (where required) were blinded to the genotype of the mice.

***Open field test (OFT)***

The OFT was administered as reported previously[3, 5], with minor alterations. Briefly, locomotor activity in an open field (W x D x H = 40 x 40 x 27 cm) was recorded for 60 min. The total distance traveled (cm) and time in center (center 2/3 of the field) from the first 5 min (sec) was analyzed to assess novelty-induced locomotor behavior. The total distance travelled during the entire 60 min session was analyzed to assess general locomotor activity.

***Elevated plus maze (EPM) Test***

The EPM was administered as reported previously[3, 5]. Briefly, the EPM consisted of two open arms (30 x 7 cm) and two closed arms (30 x 7 cm) emanating from a common central platform (7 x 7 cm), elevated to a height of 30 cm above floor level. Testing commenced by placing a mouse in the center of the maze facing an open arm, and the mouse was allowed to roam freely for 15 min. Mice were removed from analysis if they fell off the maze. The total distance travelled, entries into and time spent in open arms was analyzed.

***Y-maze test***

The Y-maze test was administered as reported previously.[3–5] Each mouse was placed in the center of a symmetrical Y maze (arm length:42 cm, wall height:15cm) and was allowed to explore freely for 10 min. Total distance traveled and arm entries was analyzed. Successful alternation rates amongst the 3 arms were used as a measure of short term spatial working memory and were calculated using a previously reported R script[4] from the arm entry data.

***Social Interaction Test***

The social interaction test[6] was administered by placing two mice into an open field (W x D x H = 40 x 40 x 27 cm) together. For tracking, mice were anesthetized with isoflurane two days before the experiment, the hair on the dorsal regions of each mouse was clipped with animal hair clippers (Panasonic), and the skin was marked with mouse-skin safe markers (Brain Science Idea) so that two mice introduced into the chamber at the same time will not have the same color. At the time of testing, two mice of the same genotype housed in different cages were introduced into the open field and allowed to interact freely for 10 minutes. Measurement was initiated after both mice were introduced into the open field. As a measure of social behavior, the number and duration (s) of contacts between the nose of each mouse and the nose, body center, and tail base of the other mouse was analyzed.

***Inhibitory avoidance test***

The inhibitory avoidance test was performed as previously described[3, 5] with minor modifications. The step-through inhibitory avoidance apparatus consisted of a straight alley divided into a light chamber (8 cm long; grey Plexiglas) illuminated by a lamp, a dark chamber (16 cm long; black plexiglas) covered by black cloth and fitted with an electric grid floor, and a black sliding door separating the two chambers. On the training day, mice were placed in the light chamber with the door leading to the dark chamber raised. Once the hindlimbs of the mice had stepped through into the dark chamber, the door was closed and an electric footshock (0.6 mA, 60 Hz, 1 s) was delivered. Retention was tested 24 hr later following a similar procedure, except no shock was delivered. On both days, latency to enter the dark chamber was measured manually with a stopwatch.

***Prepulse inhibition test***

The startle response and prepulse inhibition (PPI) to a 120 dB auditory stimulus were measured using a startle reflex measurement system (SR-LAB) as previously described.[3, 5] Test sessions consisted of six trial types (pulse-only trial, and 74, 78, 82, 86, or 90 dB prepulse trials). Seven blocks of the six trial types were presented in a pseudorandomized order such that each trial type was presented once within a block. The following formula was used to calculate PPI. 100−((Response on acoustic prepulse−pulse stimulus trials/Startle response on pulse-only trials) x 100).

**Blood and Tissue Sample Collection**

**Dissection**

Blood and brain samples were collected in the morning (0900-1130). For mice subjected to the behavioral test battery, dissection was performed at least 5 days after completion of the behavioral test battery, Mice were deeply anesthetized with isoflurane, and blood was collected from the inferior vena cava using a 26G needle and syringe. After blood collection, the mice were promptly decapitated, and the brain was removed from the cranium using tweezers.

**Collection of plasma**

The collected blood was immediately transferred to a 1.5 ml Protein LoBind Tube (Eppendorf) containing 10 µl of sodium heparin (1000 U/10 ml, Mochida Pharmaceutical) to prevent coagulation and mixed thoroughly. Whole blood was centrifuged at 1000 × g for 10 minutes at 4°C, to separate the plasma. The plasma was collected and stored at -80°C.

**Punch collection from brain samples**

The brain was immediately washed with phosphate buffered saline (PBS) in a 10-cm glass dish to wash out blood, and then sliced into 1mm thick coronal slices using a brain matrix (Brain Science Idea). 5 regions (Fig S4) were isolated from coronal slices using a razor blade and biopsy punch (1.0 mm Kai Industries), collected in 1.5 ml tubes, immediately frozen on dry ice, and stored at -80°C until further use. All procedures were done on ice to minimize sample degradation.

**Multiplex Immunoarray**

Plasma levels of cytokines, chemokines, receptors, and other immune-related signal proteins were measured using a Mouse Magnetic Luminex Assay (R&D systems), as previously reported[5]. Twenty-nine different cytokines were measured: IL-1 β, IL-2, IL-4, IL-5, IL-6, IL-7, IL-10, IL-13, IL-16, IL-17A, IL-17E, IL-33, IL-6 Rα, CCL3, CCL5, CCL11, CXCL 1, CXCL10, LIX, IFN-g, TNF-α, TNF RI, TNF RII, VEGF, PDGF-BB, Prolactin, TIMP-1, G-CSF, and GM-CSF.

**Quantitative Real-time PCR (qRT-PCR)**

Total RNA was isolated from ether tissue or cell samples using ReliaPrep™ RNA Tissue Miniprep (Promega) according to the manufacturer's specifications. Total RNA was used to perform first strand cDNA synthesis with the PrimeScript RT reagent kit (Takara Bio., Shiga, Japan). Primers used for qRT-PCR are listed in Table S19. PCR was performed on the QuantStudio 6 Flex Real-Time PCR System (Life Technologies) using GeneAce SYBR qRCR Mix α Low ROX (Nippon Gene Co., Toyama, Japan). PCR condition is as follows: Denaturation at 95 °C for 10 min; 45 cycles of amplification at 95 °C for 15 s and 60 °C for 30 s. Each sample was measured in technical triplicate, and all data are normalized to β-Actin. For expression analysis from brain tissue, data was collected using at least three independent litters of mice.

**Antibodies**

The following primary antibodies were used in the present study: NF-κB p65 (#8242; Cell Signaling Technology (CST) Inc., MA, USA), RCC1 (sc-1161; Santa Cruz, TX, USA), TDP-43 (10782-2-AP; Proteintech, IL, USA ), STAT3 (#8768; CST), MeCP2 (#3456; CST), importin α1 (KPNA1, 2A4-1B5; Abnova, Taiwan), importin α2 (KPNA2, ab84440; Abcam, MA, USA), importin α3 (KPNA3, IMG3568; Imgenex, CA, USA), importin α4 (KPNA4, ab6039; Abcam), importin α6 (KPNA6, 3F8; rat mAb[7]), importin β1 (KPNB1, sc-1919; Santa Cruz), Actin (sc-

1615; Santa Cruz, CA, USA). The secondary antibodies used for an immunofluorescence analysis were: anti-rabbit Alexa Fluor Plus 488 (A32731, Thermo Fisher Scientific Inc., Rockford, IL, USA), and anti-goat Alexa Fluor 488 (A11055) for an immunofluorescence analysis, and horseradish peroxidase (HRP)-conjugated anti-goat, anti-rabbit, or anti-mouse secondary antibodies (Jackson ImmunoResearch Inc. West Grove, PA, USA) for western blotting.

**Western blotting for brain tissues**

Approximately 20-30 mg of tissue dissected from each mouse were placed in RIPA buffer (50 mM Tris-HCl (pH7.0), 150 mM NaCl, 1 mM EDTA, 0.5% Tween, 0.1% SDS, 1% Nonidet P-40) on ice and homogenized using a 23G and 26G needle 10 times each. Then, samples were sonicated for 5 mins using an Ultrasonicator US-100 (TAITEC, Saitama, Japan) followed by centrifugation at 10,000 xg 4 ºC for 10 min to remove insoluble material. The protein concentration of the lysates were measured using a BCA protein assay kit (PIERCE, IL, USA).

Western blotting was performed with lysates from brain tissue samples separated on a 10% sodium dodecylsulfate-polyacrylamide gel electrophoresis (SDS-PAGE) gel (SuperSep™ Ace; Wako Pure Chemical Industries, Ltd., Osaka, Japan). Following transfer onto an Immobilon-P membrane (Merck Millipore, Darmstadt, Germany) using a semidry-type blotting apparatus (ATTO, Tokyo, Japan), the membrane was blocked with 3% skim milk in tris-buffered saline (TBS; 20 mM Tris-HCl (pH 7.5), 150 mM NaCl) with 0.1% Tween-20 (TBST) for 1 hr and probed with primary antibodies diluted in Can Get Signal Immunoreaction Enhancer Solution 1 (CGS1; TOYOBO, Osaka, Japan). After washing the membrane with TBST, the membrane was incubated horseradish peroxide (HRP)-conjugated secondary antibodies diluted in Can Get Signal Immunoreaction Enhancer Solution 2 (TOYOBO), and then signals were detected with Chemi-Lumi One Super reagent (nacalai tesque, Kyoto, Japan).

**Establishment of primary astrocyte and MEFs**

Brains were isolated from pups of postnatal day 1-2 and kept in 6-well dishes on ice with DMEM/F12 medium (ThermoFisher) (no serum). The olfactory bulb, cerebellum, and meninges were removed under a stereo microscope, and remaining brain tissue was minced and suspended in 3 mL of DMEM/F12 and transferred to a 15 mL falcon tube. Minced tissue was centrifuged at 1000 rpm for 5 min, the supernatant was removed, and the tissue was incubated with 5 mL of 0.25% Trypsin (nacalai tesque) with 1 µg/mL DNase I (D5025, Sigma-Aldrich) at 37 ºC for 30 min. The minced tissue suspension was passed through a 40 μm Cell Strainer (#352340, BD Falcon) with additional 5 mL of DMEM/F12. The cell suspension was centrifuged at 1000 rpm for 5 min, and then the remaining cells were suspended with 5 mL of Glia culture medium (DMEM/F12, 10% FCS, Penicillin/streptomycin (P/S)). The mixed culture of glia cells was placed in a T25 flask coated with 50 μg/mL Poly-D-lysine (PDL, P7405 or P7280, Sigma-Aldrich) and cultured in a humidified incubator (37 ºC, 5% $CO_2$) for 10-14 days. Media was changed every 3 days after initial seeding. Upon confluence in the T25 flask, the flask was shaken for 16-18 h at 120-150 rpm. Cells attached to the flask after shaking was further cultured as astrocytes.

MEFs were established as described below; for the dissection of E12.5-E13.5 mouse embryos, sterilized tweezers and scissors were used to remove the head, limbs, and internal organs in PBS (-). After the tissue was incubated with 0.025% Trypsin/EDTA at 37°C for 30 min, it was mechanically dissociated by passing through a 23 G syringe at 5-6 times, and then filtered through a 40 μm mesh into a 50 mL tube, combined with DMEM supplemented with 10% FCS, centrifuged at 1000 rpm for 5 min. The cell pellets were resuspended in DMEM with 10% FCS, and then seeded into T25 flasks, incubated until subconfluent growth.

**Indirect immunofluorescence**

An indirect immunofluorescence for primary cells was performed as described previously.[8] For observation of the p65 nuclear localization (Fig. 3A), cells were treated with TNF-α (final conc. was 20 ng/mL; Miltenyi Biotec, Bergisch Gladbach, Germany) for 30 min. Following fixation with 3.7% formaldehyde in PBS for 15 min, cells were treated with 0.1% Triton X-100 in PBS for 5 min and then blocked in 3% skim milk in PBS for Astrocytes or in blocking one Histo (Nacalai Tesque, Japan) for MEFs. Primary antibodies for p65 were incubated in appropriately diluted (1: 200 dilution) in 3% skim milk in PBS for overnight at 4ºC, and antibodies for RCC1, TDP-43, STAT3, and MeCP2 were incubated in blocking one Histo (1:20 dilution) in 0.1% Tween in PBS for overnight at 4ºC, and then following by incubated with Alexa Fluor 488 -conjugated secondary antibodies (Thermo Fisher Scientific Inc., USA) diluted (1: 200 dilution) in 0.1% tween in PBS with Nacalai histo one. Nuclei were counterstained with DAPI (1: 6,000 in PBS; Dojindo Laboratories, Kumamoto, Japan) for 30 min. The samples were examined using a confocal microscope (Leica TCS SP8).

**Immunohistochemistry on brain sections**

Immunohistochemistry with HRP was performed as described previously using Bouin's-fixed adult mouse brain tissue.[9] The experiments were repeated at least twice on two different samples, with qualitatively identical results.

**LPS Treatment of Mice used for FACS and PEC experiments**

20 hours before dissection, 1mg,/kg of LPS (L2880; Sigma-Aldrich, St. Louis, MO, USA) was intraperitoneally injected into 8 to 9 week-old WT and KO mice as an inflammatory stimulus. Animals were deeply anesthetized with isoflurane and used for subsequent cell collection and FACS experiments.

**Collection of peritoneal macrophages (PECS)**

6ml of PBS(-) was injected into the intraperitoneal space and the belly was massaged well for 30 sec. A 5ml syringe with a 24G needle was inserted intraperitoneally and approximately 5ml of peritoneal fluid was collected. Collected fluid was centrifuged for 5 min and cells were collected as peritoneal macrophages.

**MACS and FACS isolation of glial cell populations**

Adult mouse whole brain was minced, digested, and processed into cell suspensions using an Adult Brain Dissociation Kit and gentleMACS™ Octo Dissociator with Heaters (both Miltenyi Biotec). Glial cell populations (Microglia, Astrocytes) were sorted from cell suspension either by MACS (magnetic cell sorting; isolated cells used for RT-qPCR) or FACS (florescence activated cell sorting; isolated cells used for RNAseq). For MACS sorting, cells were labeled with either CD11b or ASCA2 microbeads and separated using a OctoMACS™ Separator with MS Columns (all Miltenyi Biotec). For FACS sorting, cells were stained with anti-CD45-APCvio770, anti-ACSA2-APC, anti-CD11b-vioBright-FITC, anti-O4-PE, anti-CD31-PE antibodies (all from Miltenyi Biotec) as well as DAPI. Microglia and Astrocyte populations were separated in a MACSQuant Tyto cell sorter (Miltenyi Biotec), with sort gates selecting for $CD11b^{high}$, $ASCA2^{low}$, $O4^{low}$, $CD45^{low}$, $CD11^{low}$, $CD31^{low}$ cells for Microglia and $CD11b^{low}$, $ASCA2^{high}$, $O4^{low}$, $CD45^{low}$, $CD11^{low}$, $CD31^{low}$ cells for Astrocytes. Isolated cells were measured with a MACSQuant Analyzer 10 flow cytometer (Miltenyi Biotec) to confirm purity. All samples used in RNAseq analysis were confirmed to be 90%< microglia or astrocytes (Fig S6).

**RNAseq**

Total RNA was extracted from cells with an miRNeasy Mini kit (Qiagen) according to the manufacturer's instructions. Library preparation and sequencing was performed at the NGS core facility at the Research Institute for Microbial Diseases of Osaka University. Full-length cDNA was generated using a SMART-Seq HT Kit (Takara Bio) according to the manufacturer's instructions. An Illumina library was prepared using a Nextera DNA Library Preparation Kit (Illumina) according to SMARTer kit instructions. Sequencing was performed on an Illumina NovaSeq 6000 sequencer (Illumina) in the 101+101 base paired-end mode.

Sequenced reads were mapped to the mouse reference genome (NCBI-RefSeq - GCF_000001635.27_GRCm39) using STAR 2.7.10b. RSEM-1.3.3 was used to calculate gene counts for each gene based on number of reads. Differential expression analysis of each gene was performed on the TCC-GUI[10]platform using TMM-Voom for calculation of differentially expressed genes (Table S3-4). Lists of gene sets with differential expression $p \leq 0.05$, $|\text{Fold Change}| \geq 2$ (Table S5-6) were analyzed in Enrichr[11–13] for enrichment analysis of transcription factors (ChEA 2022; Table S7-8, S11-12) and histone modifications (ENCODE Histone Modifications 2015; Table S9-10, S13-14). Transcription factor analysis was done with the wPGSA package[14] based on gene count lists generated from RSEM (data from this study; Table S15-16) or from raw count values in a previously reported data set (Marvaldi, et al[1] Supplemental file: aaz5875_tables1.xls; Table S17-18).

***Data Analysis and Statistics***

Statistical analyses and data visualization other than FACS and RNAseq analysis were performed using Prism 8.0 (GraphPad Software, La Jolla, CA). RNAseq analysis was done on R. Data are presented as the Mean ± SEM for bar graphs, with dots indicating individual data points. For box-whisker plots, data is presented as the median (center line), ±1.5 interquartile range (box), and minimum and maximum values (whiskers). For violin plots, data is presented as Median (solid line), First and Third Quartiles (dotted line). Statistical differences among three groups/factors or more were determined using a one-way or two-way analysis of variance (ANOVA) or Kruskal-Wallis test, followed by *post hoc* Tukey, Sidak, or Dunn tests. For two-way repeated measures ANOVA, Greenhouse-Geisser-corrected degrees of freedom were used for the main effect of the trial when there were more than 3 trials. All ANOVA and Kruskal-Wallis results are shown in the additional Statistical Analysis File. The number of individual mice used for analysis, as well as the number of excluded outliers are shown in Statistical Analysis File.

#### 3. References for supplemental material


1. Marvaldi L, Panayotis N, Alber S, Dagan SY, Okladnikov N, Koppel I, et al. Importin α3 regulates chronic pain pathways in peripheral sensory neurons. Sci New York N Y. 2020;369:842–846.

2. Miyamoto Y, Sasaki M, Miyata H, Monobe Y, Nagai M, Otani M, et al. Genetic loss of importin α4 causes abnormal sperm morphology and impacts on male fertility in mouse. Faseb J. 2020. 2020. https://doi.org/10.1096/fj.202000768rr.

3. Nomiya H, Sakurai K, Miyamoto Y, Oka M, Yoneda Y, Hikida T, et al. A Kpna1-deficient psychotropic drug-induced schizophrenia model mouse for studying gene–environment interactions. Sci Rep. 2024;14:3376.

4. Li S, Sakurai K, Ohgidani M, Kato TA, Hikida T. Ameliorative effects of Fingolimod (FTY720) on microglial activation and psychosis-related behavior in short term cuprizone exposed mice. Mol Brain. 2023;16:59.

5. Sakurai K, Itou T, Morita M, Kasahara E, Moriyama T, Macpherson T, et al. Effects of Importin α1/KPNA1 deletion and adolescent social isolation stress on psychiatric disorder-associated behaviors in mice. Plos One. 2021;16:e0258364.

6. Ikeda K, Sato A, Mizuguchi M. Social interaction test: a sensitive method for examining autism-related behavioral deficits. Protoc Exch. 2013. 2013. https://doi.org/10.1038/protex.2013.046.

7. Mizuguchi C, Moriyama T, Yoneda Y. Generation and Characterization of a Monoclonal Antibody Against Importin α7/NPI-2. Hybridoma. 2011;30:307–309.

8. Miyamoto Y, Itoh Y, Suzuki T, Tanaka T, Sakai Y, Koido M, et al. SARS-CoV-2 ORF6 disrupts nucleocytoplasmic trafficking to advance viral replication. Commun Biology. 2022;5:483.

9. Hogarth CA, Jans DA, Loveland KL. Subcellular distribution of importins correlates with germ cell maturation. Dev Dyn. 2007;236:2311–2320.

10. Su W, Sun J, Shimizu K, Kadota K. TCC-GUI: a Shiny-based application for differential expression analysis of RNA-Seq count data. BMC Res Notes. 2019;12:133.

11. Kuleshov MV, Jones MR, Rouillard AD, Fernandez NF, Duan Q, Wang Z, et al. Enrichr: a comprehensive gene set enrichment analysis web server 2016 update. Nucleic Acids Res. 2016;44:W90–W97.

12. Chen EY, Tan CM, Kou Y, Duan Q, Wang Z, Meirelles GV, et al. Enrichr: interactive and collaborative HTML5 gene list enrichment analysis tool. BMC Bioinform. 2013;14:128.

13. Xie Z, Bailey A, Kuleshov MV, Clarke DJB, Evangelista JE, Jenkins SL, et al. Gene Set Knowledge Discovery with Enrichr. Curr Protoc. 2021;1:e90.

14. Kawakami E, Nakaoka S, Ohta T, Kitano H. Weighted enrichment method for prediction of transcription regulators from transcriptome and global chromatin immunoprecipitation data. Nucleic Acids Res. 2016;44:5010–5021.
